# Transcriptome-wide mRNP condensation precedes stress granule formation and excludes new mRNAs

**DOI:** 10.1101/2024.04.15.589678

**Authors:** Hendrik Glauninger, Jared A.M. Bard, Caitlin J. Wong Hickernell, Karen M. Velez, Edo M. Airoldi, Weihan Li, Robert H. Singer, Sneha Paul, Jingyi Fei, Tobin R. Sosnick, Edward W. J. Wallace, D. Allan Drummond

## Abstract

Stress-induced mRNP condensation is conserved across eukaryotes, resulting in stress granule formation under intense stresses, yet the mRNA composition and function of these condensates remain unclear. Exposure of ribosome-free mRNA following stress is thought to cause condensation and stress granule formation through mRNA-sequence-dependent interactions, leading to disproportionate condensation of long mRNAs. Here we show that, in striking contrast, virtually all mRNAs condense in response to multiple stresses in budding yeast with minor length dependence and often without stress granule formation. New transcripts escape mRNP condensation, enabling their selective translation. Inhibiting translation initiation causes formation of mRNP condensates that are distinct from stress granules and P-bodies; these translation-initiation-inhibited condensates (TIICs) are omnipresent, even in unstressed cells. Stress-induced mRNAs are excluded from TIICs due to the timing of their expression, indicating determinants of escape that are independent of sequence. Together, our results reveal a previously undetected level of translation-linked molecular organization and stress-responsive regulation.

## Introduction

Cells must respond to changing environments to survive and thrive. When faced with a broad range of sudden maladaptive environmental changes—stresses—eukaryotic cells downregulate translation, induce stress-responsive transcriptional programs, and form cytosolic clusters of mRNA and proteins. When microscopically visible as foci colocalized with markers such as poly(A)-binding protein, these clusters are called stress granules (SGs)^1–7^; they coexist with other cytosolic structures including P bodies (PBs), which also accumulate mRNAs and distinct marker proteins. Stress granules are conserved across eukaryotes, and are complex examples of biomolecular condensates, membraneless structures without defined stoichiometry which form by a range of processes and which concentrate specific types of biomolecules.^8,9^ Their function remains unclear, as does the relationship between stress granule formation and the accompanying transcriptional and translational responses.^10^

Early work in multiple systems established that what are now recognized as stress granules recruit multiple RNA-binding proteins along with pre-stress mRNA, yet exclude nascent mRNA produced during stress.^11,12^ In mammalian cells, SGs were shown to nonspecifically recruit untranslated mRNA but exclude two specific stress-induced heat shock protein mRNAs, HSP70 and HSP90.^13,14^ This matched prior work on heat shock granules in plants, which recruited mRNAs encoding housekeeping proteins but not those encoding newly synthesized heat shock proteins.^6^ In glucose-starved yeast cells, induced mRNAs show complex behavior dependent on their promoter, where some induced transcripts show reduced translation and accumulate in SGs or PBs while others are soluble and translated.^15^ More recent work on glucose starvation finds most translationally repressed mRNAs outside PBs and a strong correlation between transcription and translation upon stress.^16^

Stress-triggered inhibition of translation initiation plays a central role in SG formation.^12^ Subsequent ribosome run-off, polysome disassembly, and the exposure of ribosome-free mRNA, has been proposed to serve as a template or “universal trigger” for SG assembly.^12,17–19^ Consistent with the ribosome-free RNA template model, inhibitors of translation elongation which lock ribosomes on transcripts, such as cycloheximide (CHX) and emetine, inhibit SG formation, whereas an elongation inhibitor which causes ribosome release, puromycin, promotes SG formation.^17,20^

Correlations between RNA length and SG transcriptomes have been interpreted as supporting a central role of ribosome-free RNA in stress granule formation. Measurements of the mRNA enriched in stress granules in both yeast and mammalian cells claimed that mRNA length is the main determinant of enrichment with long mRNAs accumulating in SGs, while short mRNAs are excluded.^4,21–23^ Increasing RNA length promotes RNA/protein phase separation *in vitro* by the stress-granule hub protein G3BP1,^24,25^ and single-molecule studies show that mRNA length correlates with the dwell time of mRNAs on stress granules and other condensed structures.^26^ These results fit a model in which long RNAs provide more opportunities for multivalent interactions necessary to form condensates. Yet these transcriptome-scale findings are in apparent conflict with results showing selective exclusion of stress-induced mRNAs from stress granules.^10^

In contrast, other work highlights a central role for protein components in mRNA-protein condensation and SG formation.^5,25,27–32^ In mammalian cells, the protein-protein interaction network mediated by G3BP1/2 is critical for stress granule assembly following arsenite treatment.^28^ Meanwhile blocking visible SG formation with cycloheximide does not block *in vivo* condensation of Pab1^33^ and mild stresses trigger protein condensation without SG formation.^33,34^ Indeed, multiple RNA-binding proteins, including SG markers, autonomously condense *in vivo* and *in vitro* in response to physiological stress conditions.^7,33,35–37^ This supports a model in which stages of protein condensation occur regardless of whether visible stress granules eventually appear.^10^ Whether a staged model applies to mRNA condensation remains to be studied.

Here, using a wide range of methods including biochemical fractionation by sedimentation and RNA sequencing (Sed-seq), we show that virtually all pre-stress transcripts condense during stress regardless of their lengths, even in the absence of visible stress granules. At the transcriptome scale, stress-induced transcripts escape condensates and are robustly translated. A simple explanation rationalizes stress-specific differences in condensed mRNA: pre-existing transcripts condense, and newly produced transcripts escape condensation, permitting their preferential translation. We discover that specific endogenous transcripts are condensed prior to stress, only to be released from condensates to be translated during stress. Most surprisingly, condensation is pervasive even in unstressed cells and results from inefficient translation initiation. These translation-initiation-inhibited condensates (TIICs) contain both mRNA and protein, are distinct from stress granules, and potentiate SG formation. Together, these results show that mRNA-protein condensation occurs even basally outside of stress and is measurable before visible stress granules form, expanding the importance of understanding mRNA-protein condensation for cellular physiology in and outside of stress.

## Results

### Sed-seq enables measurement of transcriptome-scale mRNA condensation

We previously used biochemical fractionation via sedimentation to isolate stress-induced proteins in condensates during heat shock in budding yeast.^38,39^ The principle of the assay is that changes in particle size induced by stress or other treatments can be measured by changes in sedimentation after centrifugation. To measure mRNA in condensates, we coupled this sedimentation assay with RNA sequencing (Sed-seq) (Figure 1A). We collected and quantified transcript abundances in total, supernatant, and pellet fractions, and estimated the proportion of each gene’s transcripts in the supernatant (pSup) using a Bayesian mixture model ^38^ validated by qPCR (Figure S1A). As in previous studies, we included the magnesium chelating agent ethylenediaminetetraacetic acid (EDTA) to disassemble polysomes which would otherwise sediment along with condensates (Figure S1B).^33,40,41^ A limitation is that any magnesium-or calcium-dependent condensates will be disrupted by this method.

**Figure 1:**
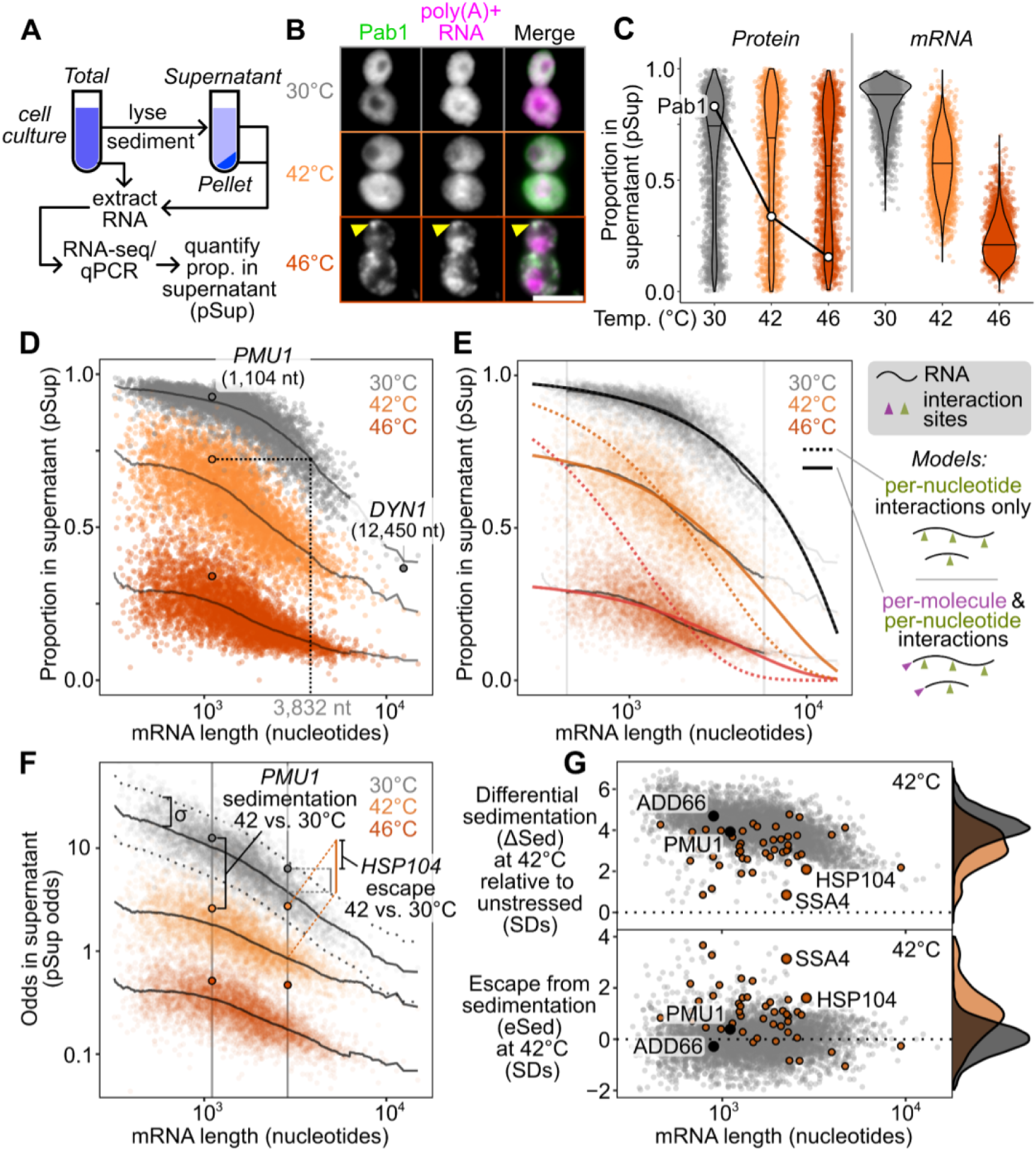
M**o**st **transcripts condense during stress, even in the absence of stress granules.** (A) Analysis of mRNA condensation by sedimentation and RNA sequencing (Sed-seq) enables calculation of mRNA proportion in the supernatant (pSup) across conditions. (B) 15 minutes of heat shock induces stress granule formation at 46°C but not at 42°C, as marked by poly(A)-binding protein (Pab1-HaloTag) and FISH against poly(A)+ RNA (scale bar = 5µm). (C) Comparison of protein condensation (data from Wallace *et al.* 2015) and mRNA condensation (this study). (D) Transcript pSup decreases with length under all conditions, including in unstressed cells at 30°C. Differences in sedimentation are caused by stress. (E) A simple clustering model (see Supplementary Text) captures average pSup and stress-induced changes. Vertical boundaries mark 1st and 99th percentile of transcripts by length. (F) Sed-seq data allow quantification of key features: differential sedimentation relative to control (fi.Sed), and escape from sedimentation (eSed). Both control for effects of transcript length and are in units of *a,* the standard deviation (SD) in sedimentation (dotted lines) around the length-dependent mean (solid lines). (G) Top, virtually all transcripts (gray points/density) sediment in response to 42°C heat shock, indicating condensation, indicated by fi.Sed>O. Bottom, most genes in the heat shock factor 1 (Hsf1) regulon (orange points/density) show significant escape from sedimentation (eSed>O).ti.Sed and eSed are in units of *a* (standard deviations, SDs).

We first used Sed-seq to examine mRNA sedimentation transcriptome-wide in unstressed conditions (30°C) and after short heat shocks at 42°C and 46°C; as expected, 46°C produced stress granules, visible as poly(A)+ RNA colocalized with foci of poly(A)-binding protein (Pab1), while the milder 42°C shock did not produce visible stress granules (Figure 1B). Sed-seq revealed large decreases in pSup across the transcriptome during heat shock, correlated with the intensity of the stress just as in the case of proteins,^33^ which we interpret as stress-induced condensation. Unlike stress-triggered protein condensation of a minority of the proteome,^38^ virtually all transcripts show substantial condensation after stress (Figure 1C). Similar to protein condensation ^7,38,42^, mRNA condensation occurs at 42°C even when SGs are not apparent. By design, Sed-seq does not enrich for mRNA association with a particular type of RNA granule, enabling an unbiased measurement of stress-induced condensation.

In our data, long transcripts showed stronger sedimentation in all conditions, including in unstressed control cells (Figure 1D), underscoring the necessity of measuring differences between treatment and control to isolate condensation. Purified total mRNA from fission yeast spiked into unstressed lysate recapitulated this length effect, and spiked-in mRNA remained soluble in stressed lysate (Figure S1C), indicating that mRNA condensation occurs before lysis and that long transcripts show increased sedimentation under all conditions, suggesting an intrinsic property—such as mass—is responsible. Indeed, a simple two-parameter physics-based model in which mRNPs sediment only due to their mass fits the average sedimentation of unstressed-cell transcripts well (Figure 1E, Figure S1D, Supplementary Text), with substantial deviation only for the longest 1% of transcripts, which sediment less than predicted.

Stress-induced condensation of RNA shows little length-dependence, and even short transcripts show a substantial increase in condensation after stress (Figure 1D). With two additional parameters reflecting stress-induced changes in the probability of inter-mRNA interactions per nucleotide (length-dependent) and per molecule (e.g. per 5′ or 3′ end, length-independent), the model fits the treatment averages closely, again deviating substantially only for the longest 1% of transcripts (Figure 1E). Length-independent interactions have stronger effects on condensation than length-dependent interactions for >99% of transcripts at 46°C (Supplementary Text), and a model without a length-independent parameter is sharply rejected for both treatments (Figure 1E, dotted lines; Figure S1D; *F* = 4134.66 (42°C), *F* = 21,507.21 (46°C), *P* < 10^−6^ in each case; Supplementary Text). These results reveal a surprisingly minor role for interactions correlated with transcript length in promoting condensation.

The systematic relationship between pSup and mRNA length places bounds on the size of stress-induced condensates (see Supplemental Text). For example, 1.1-kilobase *PMU1* transcripts sediment after 42°C heat shock as if they were at least three times their unstressed size, and after 46°C shock as if more than ten times their unstressed size, lower than the heaviest detected mRNP in unstressed yeast, the 12.4-kilobase transcript encoding dynein (*DYN1*) (Figure 1D). These several-fold size changes, regardless of length, are precisely what is expected from condensation and rule out alternative explanations such as the formation of a large fixed-size complex (e.g. a stalled initiation complex) on individual mRNPs.

We next derive an estimate of condensation per mRNA that is corrected for length-dependent baseline sedimentation. Using the log-odds pSup to prevent compression at very high or low pSup values (Figure 1F), and taking a windowed average pSup as a function of transcript length for each treatment, we then calculate a differential sedimentation score (ΔSed), the difference between treatment and control in units of σ, the standard deviation (SD) of the unstressed control around this windowed average. ΔSed quantifies the increase in sedimentation due to stress, which we interpret as condensation. Even if a transcript condenses during stress (ΔSed>0), it may do so less or more than other transcripts of the same length. To quantify the escape from differential sedimentation (eSed) we score the difference between a particular transcript’s ΔSed and the mean ΔSed of transcripts of the same length, again in units of σ (Figure 1G).

We noted that a small set of transcripts showed significantly different changes in sedimentation and escape in response to stress relative to the rest of the transcriptome. For example, genes regulated by heat shock factor 1 (Hsf1), the master regulator of the core heat-shock response, showed less sedimentation and greater escape (Figure 1G).

However, our conclusions differ substantially from another report of the stress-granule transcriptome in yeast, which concluded that transcripts accumulate in SGs in proportion to their length.^4^ In this prior study, differential sedimentation is the only means by which condensed material is enriched,^43^ justifying a direct comparison. We hypothesized that the prior study’s inability to correct for length-based sedimentation, due to lack of a non-stress control, created an artifactual enrichment for long transcripts. To match stress conditions, we treated cells with 0.5% azide or mock conditions and performed Sed-seq. We found that ΔSed in these data and previously reported SG enrichment were anticorrelated (*r*=−0.3, *P*<10^−6^) (Figure S1F). Our Sed-seq results on unstressed cells reproduce the previously reported results to a high degree of accuracy (*r*=0.77, *P* < 10^−6^, Figure S1G), which is here due to stress-independent sedimentation of long transcripts, not stress granule formation. Thus, controlling for mRNA length is necessary to avoid artifactual conclusions, and to extract signals of biological regulation from sedimentation-derived data.

### Stress-induced mRNAs escape condensation and are preferentially translated

The apparent escape of Hsf1-regulon transcripts from condensation during heat shock (Figure 1G) prompted us to ask whether stress-induced transcripts in general were more likely to escape condensation. Indeed, transcripts that are highly induced during stress strongly tend to escape condensation (Figure 2A). More specifically, genes regulated by the core heat shock response transcription factor Hsf1^44^ tend to escape condensation (eSed > 0) during heat shock (Figure S2A,B Wilcoxon rank sum test *P* < 10^−6^). Escape is not specific to Hsf1 targets, as most stress-induced genes also escape condensation, including, at 42°C, targets of Msn2/4^45^, another stress-activated transcription factor (Figure S2A,B, Wilcoxon rank sum test *P* < 10^−6^). The degree of induction correlated with the degree of escape, suggesting that pre-stress mRNAs even from stress regulons condensed, and thus hinting that regulation by specific transcription factors was not the primary determinant of escape.

**Figure 2:**
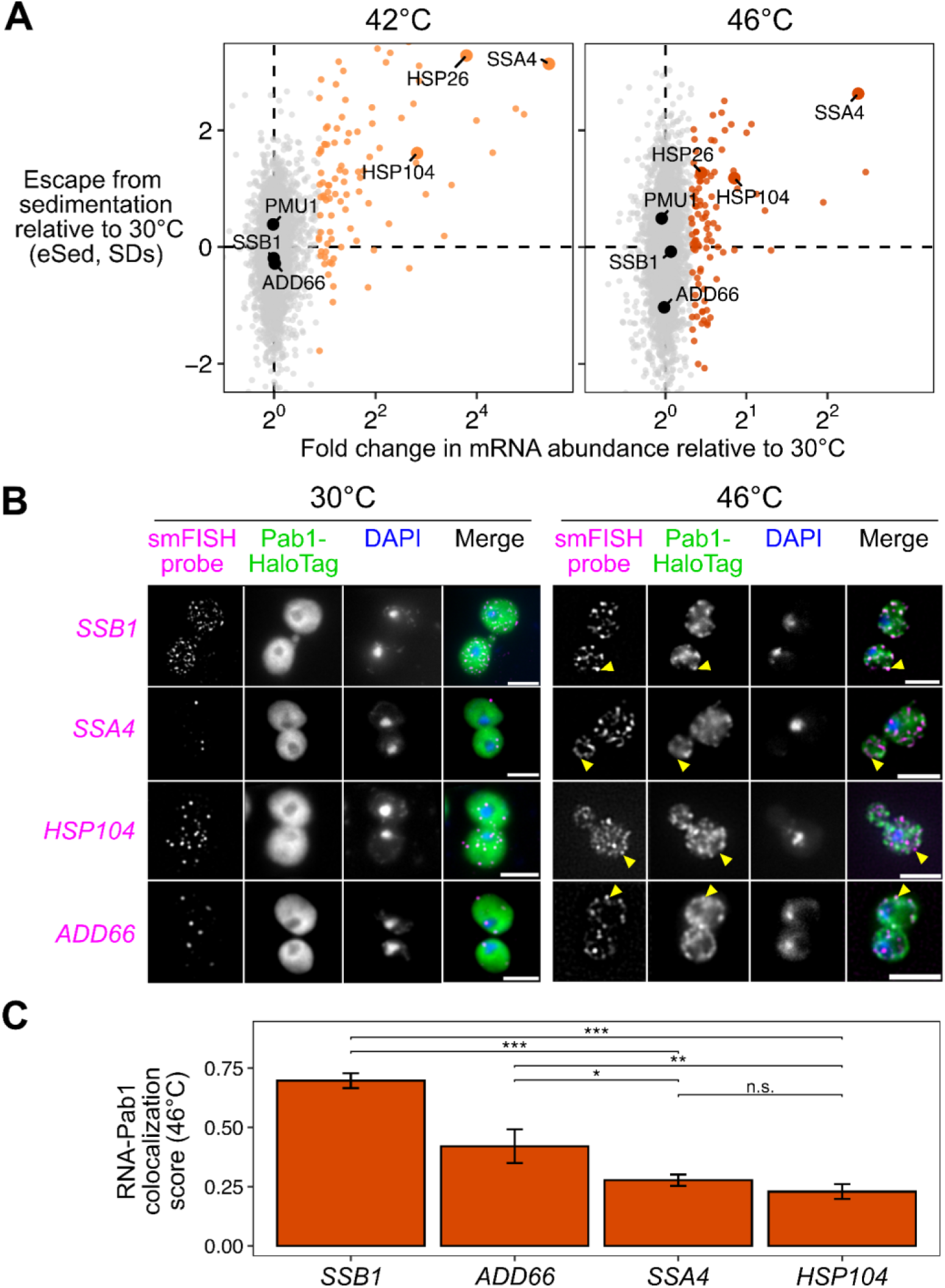
I**n**duced **transcripts escape condensation during heat shock. (A)** Comparison of mRNA abundance changes during heat shock reveals that transcript induction quantitatively predicts escape from condensation. In color are the top 100 most induced genes for each respective stress treatment. Labeled are genes which are mentioned elsewhere in the manuscript. (B) smFISH of induced *(SSA4/HSP104)* and uninduced *(SSB1/ADD66)* transcripts confirms that induced mRNA are not localized to Pab1-HaloTag marked stress granules. Scale bars are 5 µm. (C) Colocalization was quantified by comparing the intensity of the Pab1 channel in regions with mRNA foci to random regions in each cell. The colocalization score is plotted as the mean of all cells in each condition. Pairwise Welch’s t-tests were performed P-values were adjusted using the Holm method to correct for multiple comparisons. Significance thresholds were defined as follows: N.S. (p 2! 0.05); * (p < 0.05); ** (p < 0.01); *** (p < 0.001)

Stress-induced transcripts escape condensation even under conditions without apparent stress granules (e.g. 42°C). Are they also excluded from stress granules? To answer this question, we used single-molecule fluorescence in situ hybridization (smFISH)^46^ to examine the relative localization of transcripts to stress granules. We initially focused on two transcripts of nearly identical length, both encoding Hsp70 chaperones: *SSB1/2* transcripts, encoding a cytosolic Hsp70 species which is abundant in unstressed cells, and *SSA4* transcripts, encoding a stress-induced cytosolic Hsp70. We predicted that the induced *SSA4* transcripts would be excluded from stress granules. Consistent with our Sed-seq results, in 46°C heat-shocked cells, *SSB1/2* transcripts colocalized with stress granules marked by Pab1, while *SSA4* transcripts were largely excluded (Figure 2B). We then picked another pair of transcripts to test a key observation from our Sed-seq data: that length was not a determining factor in stress granule recruitment or exclusion. Indeed, induced long (2873 nt) *HSP104* transcripts were excluded, and uninduced short (896 nt) *ADD66* transcripts were recruited (Figure 2B). In order to quantify this observation, we calculated the intensity of the Pab1 channel in regions with mRNA and compared that to random regions around each cell (Methods). Reflecting the extent of the colocalization between the mRNAs and stress granules, *SSB1* and *ADD66* containing regions are strongly enriched for Pab1 signal upon stress, while *SSA4* and *HSP104* are only slightly enriched (Figure 2C, S2C). Together, Sed-seq and smFISH results form a consistent picture in which, regardless of length, stress-induced transcripts disproportionately escape stress-induced mRNP condensation.

Is the escape of induced transcripts from condensation specific to heat shock? To answer this question, we carried out Sed-seq on cells exposed to different stresses known to trigger stress granules: sodium azide (NaN_3_)^4,43,47,48^ or ethanol^49^ (Figure 3A). Following previous literature, we tracked SG formation using Pab1-GFP for heat shock and NaN_3_ stress, and Pbp1-GFP for ethanol stress.^38,47,49^ Across three types of stress, only severe stress triggered visible granule formation, while transcriptome-wide mRNA condensation occurred for all studied stress levels (Figure 3B). We find little evidence for increased stress-induced condensation of long transcripts for any of these stresses (Figure S3A). The magnitude and variability of condensation varied across these diverse stresses but followed a consistent dose-dependent pattern.

**Figure 3:**
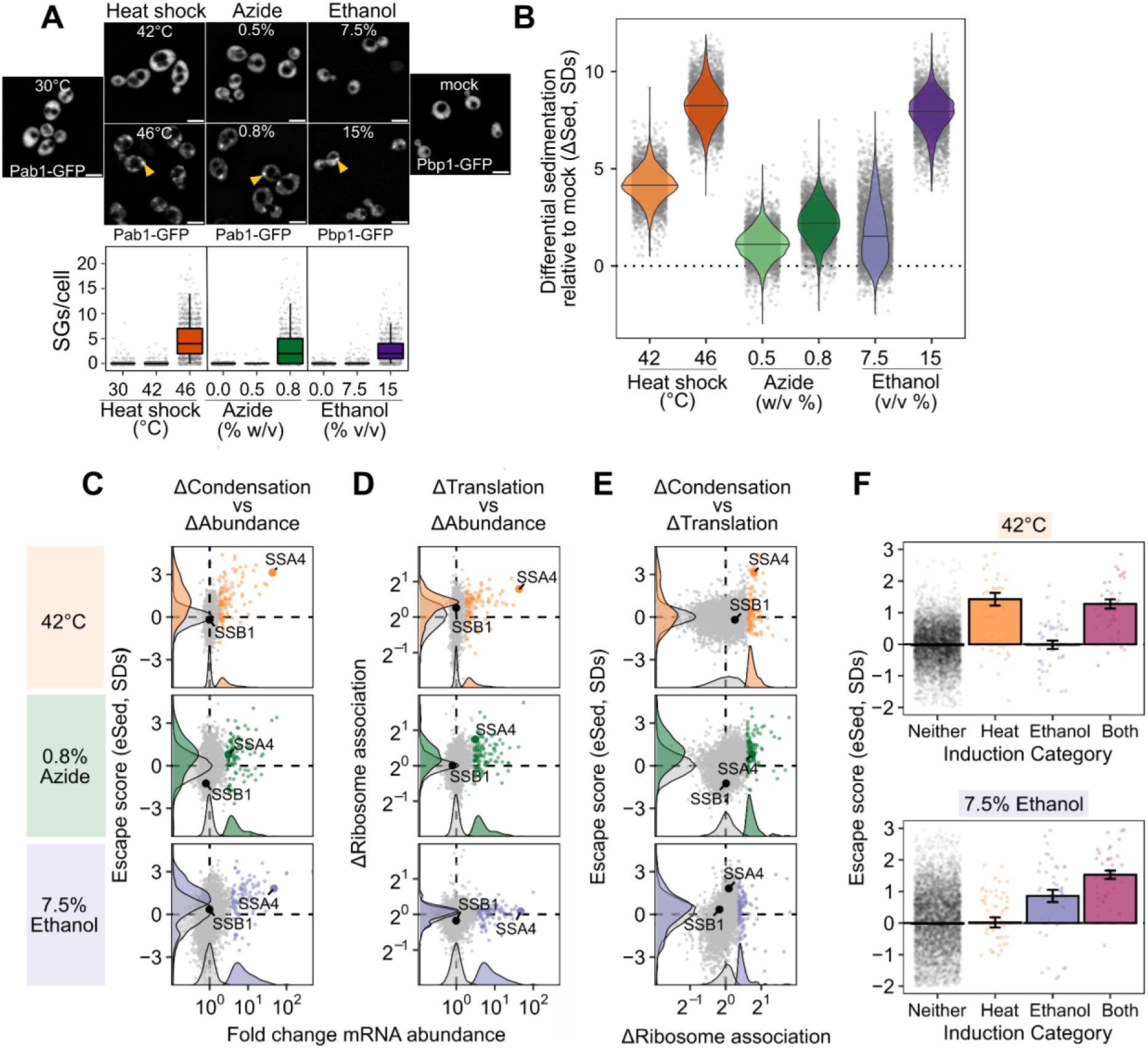
Newly transcribed and well-translated mRNAs escape condensation across stresses. (A) Severe, but not mild, stress induces visible SGs across multiple conditions. Scale bar is 5 ym. (B) Both mild and severe stress induce transcriptome-wide sedimentation of mRNA, with the extent of pelleting correlating with the severity of the stress. (C) Across stresses, the most induced mRNA (top 100 induced transcripts are highlighted) escape from condensation. (D) Polysome-seq was used to measure the stress-induced change in ribosome association (top 100 induced transcripts are highlighted). (E) Directly comparing changes in translation and sedimentation (top 100 translationally upregulated transcripts are highlighted) shows that well-translated messages during stress tend to escape condensation. (F) Transcripts from genes induced during heat shock, but not ethanol stress, also escape condensation during heat shock, but not ethanol stress, and vice versa.

Strikingly, stress-induced transcripts relatively escaped condensation across all three stresses (Figure 3C, S3C) as quantified by eSed. This result echoes early results reporting exclusion of nascent transcripts from SGs.^11,12^ In contrast, induced transcripts are not depleted from the previously reported SG transcriptome (Figure S3B).^4^

Do the different transcripts escape mRNA condensation in response to different stresses? Comparison of the eSed scores between stresses addresses this question. Comparing the transcripts which are specifically induced during heat shock, ethanol stress, both, or neither, finding that transcripts escape condensation if only when induced in that specific stress (Figure 3F).

To what extent does mRNA translation correlate with escape from condensation? We measured mRNA-ribosome association transcriptome-wide by isolating and sequencing mRNA from polysome gradients, quantifying the stress-induced change in ribosome association on each transcript (Polysome-seq).^50^ In heat and azide stress, but not ethanol stress, induced transcripts tended to be preferentially translated (Figure 3D, S3D). Preferentially translated transcripts tend to escape condensation in all stresses (Figure 3E, S3E). Transcriptional induction, escape from condensation, and increased translation co-vary in each stress condition, suggesting a functional role for condensation in translational repression of pre-existing transcripts. To establish causality, we turned to synthetic reporter constructs.

### Transcript age and translation independently regulate condensation during stress

The observation that stress-induced transcripts escape condensation is consistent with a model in which newly produced transcripts are protected from condensation for some time during stress, regardless of their identity. This new-transcript model predicts transcript exclusion will correlate with the level of induction, which is directly related to the proportion of transcripts which are new during stress, assuming degradation can be neglected. A major alternative to the new-transcript model is that sequence-encoded mRNA features, such as structure or the presence of specific motifs or untranslated-region (UTR) binding sites, determine escape. This alternative model predicts that transcripts will escape condensation independent of induction level. Sed-seq data are consistent with the new-transcript model, showing escape from condensation strongly depends on induction level (Figure 3C, S3C).

If timing of transcript production largely drives escape from condensation, then synthetic transcripts expressed from stress-independent inducible promoters should have their condensation determined by when their expression occurs. We built TET-inducible reporters with regulatory regions (5′ and 3′ UTRs) from genes which are heat-induced (*HSP26*) and heat-insensitive (*PMU1*, whose condensation behavior follows the bulk pre-stress transcriptome) (Figure 1G) ^51^. We induced each reporter before and during heat shock, and measured its condensation behavior via sedimentation with qPCR (Figure 4A,B). Both reporters were uncondensed at 30°C, and condensed at 42°C and 46°C when expressed prior to heat shock. Both, however, showed substantially reduced condensation when newly expressed during heat shock (Figure 4A,B, ANOVA *P* < 0.001). These results demonstrate that the timing of expression is a primary determinant of a transcript’s condensation fate. Transcripts which are newly produced during stress will escape condensation to a significant degree, independent of their sequence. On the other hand, transcripts produced before stress, even if they contain the sequence of a stress-induced gene such as *HSP26*, will nevertheless condense during stress.

**Figure 4:**
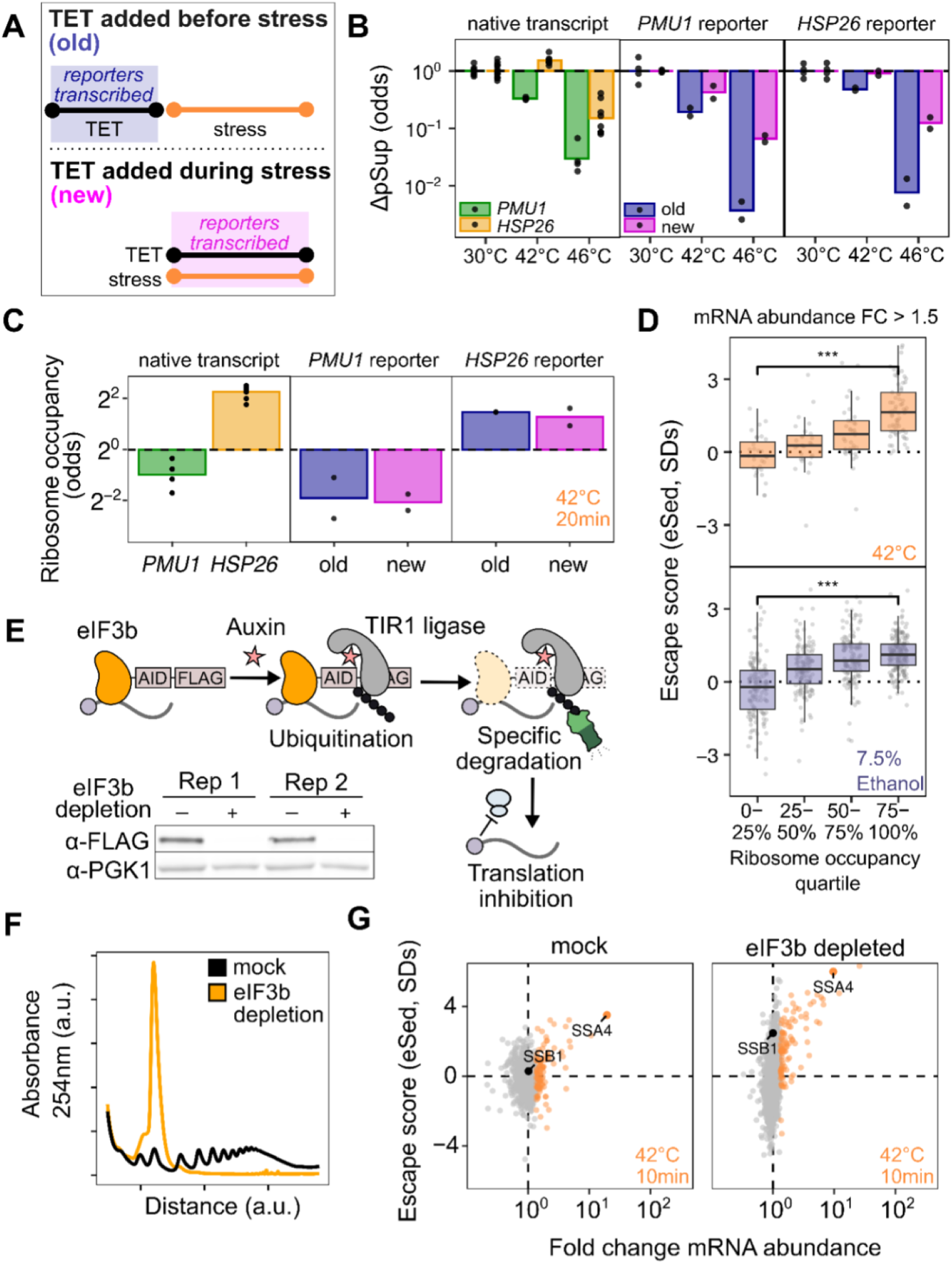
T**r**anslation **and induction are independently sufficient to promote escape from condensation.** (A) Inducible reporter transcripts with sequences derived from an induced transcript *(HSP26)* or an uninduced transcript *(PMU1).* (B) “New” transcripts sediment less than “old” transcripts for both reporters, as measured by centrifugation and qPCR after 10 minutes of stress. (C) The HSP26-derived reporter transcript is better translated than the *PMU1* reporter regardless of age, as measured by qPCR analysis of ribosome association using sucrose cushions after 20 minutes of stress. (D) Analysis of transcriptome-wide data in Figure 3 shows that even among induced mRNAs, ribosome binding is correlated with escape from condensation (Wilcoxon rank sum test, N.S.: P.:: 0.05; ••: *P* < 0.01; •••: *P* < 0.001). (E) The auxin-induced degradation system was used to deplete the translation initiation factor elF3b. (F) Depletion of elF3b leads to translational collapse as measured by polysome profiles. (G) Even in the absence of translation initiation, stress-induced transcripts still escape condensation after 10 minutes of 42°C stress (highlighted: top 100 induced transcripts per condition).

Given the clear relationship between transcript induction and escape from condensation, we sought to understand how translation fits into this model. We measured ribosome occupancy by sedimenting lysate through a sucrose cushion and quantifying the ribosome-free abundance in the supernatant and the ribosome-bound abundance in the pellet, after correcting for condensed mRNA which pellets even in EDTA buffer (Figure S4A-D)^52^. We found that, after 20 minutes of 42°C stress, the *HSP26* reporter had high levels of ribosome occupancy while the *PMU1* reporter had low ribosome occupancy regardless of whether the transcripts were new or old (Figure 4C). This translational difference matched the behavior of the native transcripts; native *HSP26* transcripts have a higher ribosome occupancy than native *PMU1* transcripts across conditions. These results show that the escape from condensation of new transcripts is not a simple consequence of their translation status.

While the reporters show that condensation can be altered independently of translation, the poorly translated *PMU1* reporters do condense more than the *HSP26* reporters (ANOVA *P* < 0.001). This is reflected in the transcriptome-wide data, which show that even amongst transcriptionally induced transcripts, poorly translated mRNAs do not escape from condensation (Figure 4D). To further investigate the relationship between translation and condensation, we generated a strain of yeast with an auxin-inducible degron (AID) tag on the C-terminus of eIF3b, a subunit of the essential initiation factor eIF3 ^53,54^. Western blotting confirmed successful degradation (Figure 4E), which resulted in polysome collapse (Figure 4F). We then performed Sed-seq on samples heat-shocked after two hours of mock treatment or eIF3b depletion. Even in cells with translation initiation blocked by eIF3b depletion, induced messages escape condensation (Figure 4G), indicating that active translation is not required for escape. Together, these results indicate that two factors simultaneously contribute to escape from condensation: being newly transcribed during stress and being well-translated.

### Translation-inhibition-induced condensates (TIICs) of mRNA and protein precede stress granule formation and form in the absence of stress

What causes condensation? The inhibition of translation initiation plays a central role in most models, through the resulting ribosome-free mRNA that has been thought to mediate condensation ^18^. Differences in translation initiation are present in cells even in the absence of stress, raising the possibility that condensation occurs under a wide range of conditions. We therefore first looked at the relationship between relative sedimentation and translation. We quantified the sedimentation of each transcripts within a condition relative to the mean of similar-length transcripts (rSed) again expressed in standard deviation units, σ).

In untreated cells at 30°C, ribosome occupancy and relative sedimentation were inversely correlated (*r*=−0.65, *P* < 10^−6^, Figure 5A). Transcripts of *HAC1*, encoding the master regulator of the unfolded protein response (UPR), drew our attention given their high rSed and low occupancy. *HAC1* mRNA relies on a long-range base-pairing interaction between its 5′ UTR and unspliced intron to block translation initiation ^55^. In contrast, the other abundant mRNA in yeast that is translationally repressed in unstressed cells–GCN4, encoding the master regulator of the amino acid starvation response—initiates translation normally on an upstream reading frame which prevents translation of the main coding region ^56^. Both mRNAs have similar lengths (*GCN4*: 1465 nucleotides, *HAC1*: 1197 nucleotides), and both are in the bottom 5% of all transcripts for ribosome occupancy (Figure 5A). Yet while *GCN4* has a rSed near the mean (rSed=−0.08, 45th percentile), *HAC1* sediments far more than the transcriptome average in unstressed cells (rSed=0.95, 95th percentile) (Figure 5A).

**Figure 5:**
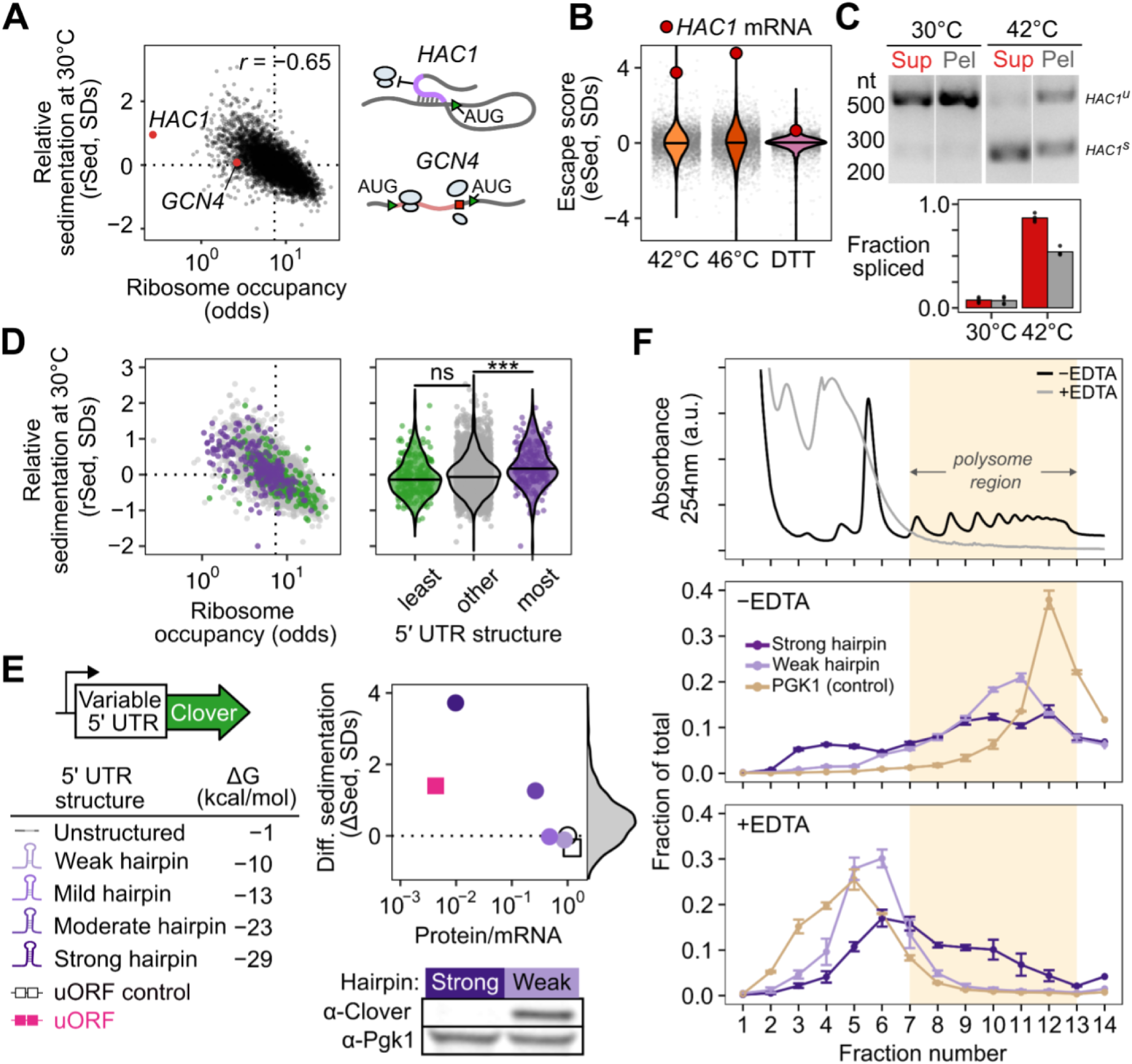
T**r**anslation**-initiation-inhibited condensates (TIICs) form in the absence of stress.** (A) Relative sedimentation (rSed) and translation measured by ribosome occupancy are negatively correlated in unstressed cells. Initiation-blocked *HAC1* shows strong sedimentation, but uORF-regulated *GCN4* does not. (B) *HAC1* mRNA becomes less condensed during heat shock and OTT treatment, as measured by escape from sedimentation. (C) 42°C treatment leads to splicing of *HAC1* mRNA as measured by RT-PCR of supernatant (red) and pellet (grey) fractions. Both the spliced *(HAC1^u^)* and unspliced *(HAC1u)* forms are present after treatment. (D) Left: Ribosome occupancy in unstressed cells correlates well with length-normalized sedimentation. Right: The amount of computationally predicted structure in the 5’ UTR of transcripts predicts their relative sedimentation. (E) Sedimentation reporters with variable 5’UTRs were generated, which repress translation via either structured hairpin or uORF sequences. Translation was quantified by the ratio of steady-state protein level to mRNA abundance and by western blot. (F) Polysomes and associated *PGK1* (control) and weak hairpin mRNAs can be disrupted with EDTA, exposing polysome-scale condensates containing initiation-blocked strong hairpin mRNA.

During heat shock at 42°C and 46°, *HAC1* mRNA showed strong escape from condensation (Figure 5B), despite showing no transcriptional induction (Figure S5A). The translation initiation inhibition of *HAC1* is relieved by mRNA splicing in the cytoplasm, leading to translation of the encoded Hac1 transcription factor, Hac1 nuclear import, and subsequent UPR activation.^55,57^ Although this process is insensitive to heat shock at 37°C,^55^ induction of *HAC1* splicing has been observed after hours of growth at 39°C.^58^ We hypothesized that more robust heat shock above 42°C caused dissolution of condensates containing *HAC1* mRNA corresponding to relief of translation initiation inhibition by splicing. Multiple predictions follow: 1) *HAC1* TIIC dissolution should occur during activation by other UPR triggers; 2) *HAC1* should be spliced in response to the short heat shocks that trigger TIIC dissolution; 3) if *HAC1* mRNA is translated, the resulting Hac1 transcription factor should drive transcription of UPR genes.

We tested each of these predictions in turn. First, we performed Sed-seq on cells treated with DTT, a standard UPR trigger. Confirming our prediction, *HAC1* mRNA showed among the strongest condensate escape across the entire transcriptome upon DTT treatment (Figure 5C) again despite showing no transcriptional induction (Figure S5A). Reductions in *HAC1* relative sedimentation accompanied increases in ribosome association across all stresses (Figure S5B). Second, we examined *HAC1* splicing in response to an 8-minute, 42°C heat shock. Before shock, *HAC1* mRNA was unspliced, running as a single large band. After shock, the spliced form of *HAC1* appeared as a smaller band (Figure 5D), confirming our second prediction. Under these conditions, *HAC1* is not completely spliced, and only the spliced form of *HAC1* partitioned into the soluble fraction (Figure 5D). These observations are consistent with release from condensates only of spliced *HAC1* transcripts in concert with their translational activation, while unspliced, initiation-blocked *HAC1* transcripts remain condensed.

Third, we looked for transcription of UPR genes at 42°C, as identified in Kimata et al., 2006.^59^ We observed a slight but unmistakable induction after a 10-minute 42°C shock (Figure S5C, Wilcoxon rank sum test *P* < 10^−6^). Next, we predicted that other heat-shock data would show induction of the UPR at 42°C, which requires that active Hac1 protein be translated. Indeed, data from a systematic study of the heat shock response in budding yeast^60^ revealed that UPR targets were significantly induced by 10-or 30-minute shocks at 42°C (Wilcoxon test *P* values < 10^−3^ in both cases), but not at 37°C (Wilcoxon test *P* = 0.15 and 0.70) (Figure S5D).

Together, these results support a simple and previously unappreciated sequence of events during *HAC1* activation: *HAC1* mRNA resides in initiation-blocked condensates under basal conditions, and is spliced and released from condensates upon UPR-inducing stress, coinciding with translation of the Hac1 transcription factor protein which then drives UPR regulon transcription.

More broadly, *HAC1* mRNA condensates appear to be an extreme example of a transcriptome-scale phenomenon linking reduction in translation with condensation (Figure 5A). Lack of condensation by uORF-regulated, freely initiating *GCN4* mRNA suggests that a block in initiation, rather than other correlates of blocked translation such as ribosome-free mRNA, is the key correlate of condensation. To further test this result, we divided transcripts by the strength of the secondary structure in their 5′ UTR, a feature known to predict the translation initiation efficiency of a transcript^61^ (Figure 5E). Transcripts with the most predicted structure in their 5′ UTR had higher rSed than the bulk transcriptome.

These results provide evidence that, transcriptome-wide, mRNAs inhibited in translation initiation are found in condensates, even in unstressed cells. Anticipating later results indicating these condensates are distinct from previously described bodies, we refer to them as translation-initiation-inhibited condensates (TIICs, pronounced “ticks”).

Does inhibition of translation initiation cause TIIC formation? We asked whether we could recapitulate *in vivo* endogenous transcript-specific condensation using a series of synthetic mRNAs encoding the green fluorescent protein Clover, with progressively stronger translation initiation blocks created by hairpins in their 5′ UTR.^62^ These hairpin series blocked translation initiation, as measured by the ratio of fluorescence intensity to mRNA abundance, with more-stable hairpins more completely blocking translation (Figure 5E). Western blotting against Clover confirmed translation was permitted by the weak hairpin and blocked by the strong hairpin (Figure 5E).

As predicted, these constructs exhibited increased sedimentation inversely correlated with their translation, mirroring *HAC1* mRNA. In contrast, a synthetic uORF construct built from the *GCN4* 5′ UTR yielded substantially less condensation than the most stable hairpin construct, despite showing stronger translational repression (Figure 5E). A control construct with five point mutations disrupting the start codon in each uORF ^63^ promoted translation of the main open reading frame, as expected, and only modestly increased transcript solubility. These experiments demonstrate that even in unstressed cells, translation initiation inhibition causes mRNA condensation, producing TIICs.

### TIICs are polysome-scale mRNP condensates

To further characterize TIICs, we performed polysome profiling, separating mRNP species by size on a sucrose gradient, followed by qPCR on strains expressing strong or weak hairpin reporters (Figure 5F). The weak hairpin reporter transcript, along with the endogenous housekeeping transcript *PGK1*, co-sedimented with the heavy polysome-associated fractions, consistent with active translation and our western blot data. Strikingly, the strong hairpin reporter transcript also co-sedimented with heavy polysome fractions despite being translationally repressed.

To determine whether strong hairpin sedimentation was due to TIIC formation rather than residual ribosome association, we dissolved ribosomes and polysomes by treating lysate with EDTA prior to sedimentation. As expected, the weak hairpin and *PGK1* transcripts shifted to lighter fractions, consistent with loss of ribosome association. In contrast, the strong hairpin transcript remained in heavy fractions, demonstrating that TIICs are resistant to EDTA treatment and sediment based on size rather than ribosomal binding (Figure 5F).

To test whether TIICs pellet in heavy fractions due to membrane association, we performed a membrane flotation assay ^64^. Lysate from the strain expressing the strong hairpin reporter was spun on an iodixanol gradient in which membrane-associated molecules float and soluble components sink (Figure S5E). The strong hairpin transcript was detected in bottom fractions regardless of the presence of membrane-dissolving Triton X-100, similar to *PGK1* and *TUB2* transcripts (Figure S5F), indicating that TIIC sedimentation is not due to membrane association. This is supported by transcriptome-wide analysis of relative sedimentation, which shows that transcripts encoding secreted proteins do not show increased sedimentation (Wilcoxon rank sum test *P* > 0.99, Figure S5G) ^65^.

These data confirm the presence of mRNP condensates triggered by translation-initiation inhibition in unstressed cells, using a method distinct from Sed-seq. The observation that TIICs display sizes comparable to polysomes provides additional insight. A polysome with five ribosomes weighs approximately 17 MDa (each yeast ribosome weighs about 3.3 MDa, and the strong hairpin reporter transcript is ∼1.8 kb, which is roughly 0.54 MDa). This size is consistent with TIICs being too small to be visible via standard microscopy, large enough to be detected by sedimentation assays, and too large to be a single mRNP. Given that the strong hairpin transcript is 1800 nucleotides long, to reach 17 MDa would require binding of one 65 kDa RNA-binding protein (e.g., Pab1) every 7 nucleotides, which is unphysical (e.g., Pab1 requires 12 nucleotides for full-affinity binding).

Together, these results further confirm the identification of TIICs as mRNP condensates, unassociated with membranes, which appear in a range of sizes up to several megadaltons.

### Blocking translation initiation at distinct steps causes global mRNA condensation and implicates an upstream, competitive step

The results above show that blocking initiation in a single reporter transcript triggers its condensation. To examine the results of a global blockade of initiation, we generated different yeast strains with auxin-inducible degron (AID) tags on eight eukaryotic initiation factors acting at multiple initiation stages (Figure 6A,B) ^53,54^. Western blotting confirmed successful translation initiation factor degradation after two hours (Figure 6C), which resulted in polysome collapse (Figure S6A) and proteome-wide reduction in translation activity (Figure 6D, Figure S6B–D).

**Figure 6:**
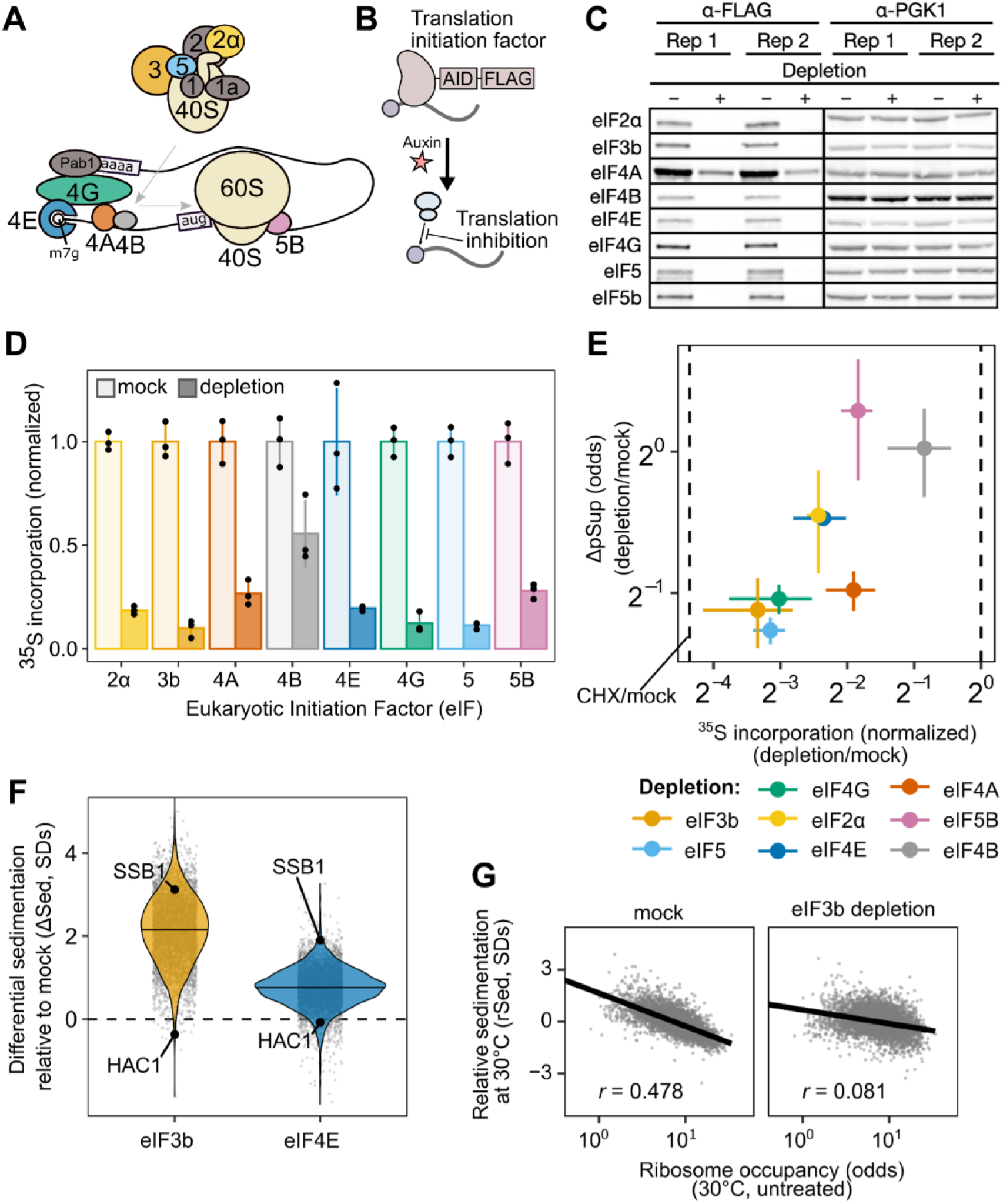
global translational initiation inhibition triggers transcriptome-wide TIICs. (A) Translation initiation factors involved in various steps of initiation were (B) depleted via the auxin-inducible degradation system. (C) Depletion for each factor was verified via western blot with Pgk1 used as a control. (D) The effect on global translation level caused by each initiation factor was tested by measuring the incorporation of radiolabeled amino acids. Each depletion caused a drop in translation to varying amounts. (E) The pSup of *PGK1* and *BEM2* transcripts (mean of the two is plotted) is strongly related to the amount of translation block caused by each initiation factor depletion, suggesting that none o1 these factors are essential for condensation. (F) Sed-seq was used after elF3b and elF4E depletion to measure global sedimentation. Depletion of both factors, and especially elF3b, triggers global condensation-TIIC formation. (G) Left: The relative sedimentation of transcripts correlates well with ribosome occupancy in the mock treated sample, but this association is attenuated after elF3b depletion (r = Pearson correlation).

We used qPCR to quantify the average pSup of two transcripts, *PGK1* and *BEM2*, following two hours of initiation factor depletion. As predicted, blocking initiation triggered mRNA condensation, with the degree of translation initiation block correlating with the extent of resulting mRNA condensation (Figure 6E). Depletion of eIF4B and eIF5B caused negligible condensation, but also had the smallest effect on translation. By contrast, eIF4A depletion caused particularly strong mRNA condensation, consistent with previous evidence showing that eIF4A inhibition can trigger SG formation ^66,67^.

During global translation initiation block, we expect that all transcripts will form TIICs, leading to increased mRNA sedimentation transcriptome-wide. To test this hypothesis, we performed Sed-seq on strains depleted for eIF4E, the mRNA cap-binding protein, and for eIF3b, the factor whose depletion led to the most severe block in translation. We observed transcriptome-scale mRNA condensation in both cases, to a profound degree after eIF3b depletion (Figure 6F). Because translationally repressed mRNAs already enter TIICs in untreated cells, we predicted that they would show the smallest differences in sedimentation. Consistent with this prediction, initiation-inhibited *HAC1* mRNA showed almost no change after both depletions, whereas initiation-competent *SSB1* mRNA had a high ΔSed score (Figure 6F). Furthermore, reflecting the global convergence of sedimentation behavior during severe initiation block, the rSed scores of transcripts in eIF3b-depleted cells are much less correlated with ribosome occupancy (Spearman *r* = 0.081) than the rSed scores of transcripts in mock-treated cells (Spearman *r* = 0.48) (Figure 6G).

Together, these results show that blocking translation initiation globally triggers global mRNP condensation and augments TIICs which are present in unstressed cells. We next sought to understand the relationship between TIICs, stress-induced mRNP condensation, and stress granules.

### TIICs are stress-granule precursors

We counted stress granules before and after inhibiting translation initiation by eIF3b depletion, both in otherwise untreated and in heat-shocked cells (Figure 7A). Because automated counting scored some unstressed (30°C) cells as having multiple SGs, and all conditions show some degree of cell-to-cell variability, we scored populations of cells as SG-negative if the median number of SGs per cell was zero, and as SG-positive otherwise. Using this threshold, unstressed cells are SG-negative and cells shocked at 46°C are SG-positive (Figure 7A).

**Figure 7:**
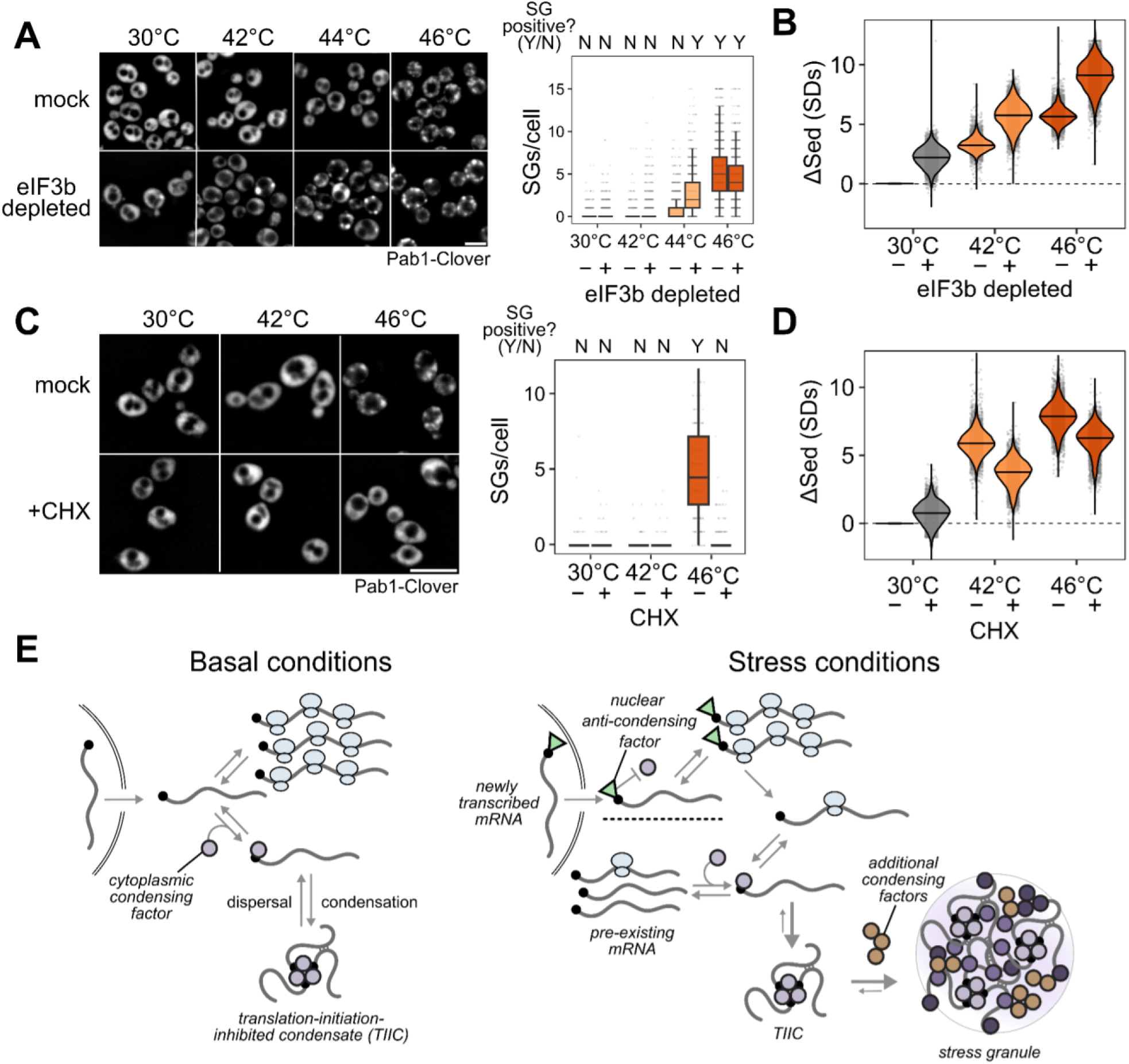
T**I**ICs **precede and potentiate stress granules.** (A) Stress granules are potentiated in elF3b-depleted cells, shown by the appearance and penetrance of stress granules at lower temperatures compared to mock-treated cells. Right: Quantification of the presence of stress granules in all conditions. (B) Sed-seq data comparing global condensation in elF3b depleted and mock cells after two hours of depletion followed by ten minutes of heat shock. elF3b depletion triggers more RNA condensation in each temperature. (C) Ten minutes of cycloheximide (CHX) treatment prior to stress prevents visible SG formation. (D) CHX treatment inhibits, but does not prevent stress-induced **RNA** condensation. (E) Model of the competition between translation initiation and TIIC formation during normal growth and stress. Well-translated transcripts are protected from condensation by competition between translation initiation and TIIC formation. During stress, newly transcribed transcripts escape stress-induced condensation, likely due to a 5’ bound protein or modification which inhibits condensation. In addition, global inhibition of translation leads to transcriptome-wide TIIC formation. These TIICs are precursors of visible stress granules, whose formation involves additional stress-induced condensing factors.

After eIF3b depletion at 30°C, which causes substantial transcriptome-wide mRNP condensation (Figure 7B), cells are SG-negative (Figure 7A). We conclude that inhibiting translation initiation by eIF3b depletion causes TIIC formation but not SG formation.

Upon heat shock at 44°C, otherwise untreated cells are SG-negative, but when eIF3b is depleted, cells become SG-positive (Figure 7A). Thus, eIF3b depletion potentiates SG formation, suggesting that TIICs are the building blocks for stress granules.

In every case, heat stress amplifies the sedimentation induced by translation initiation depletion (Figure 7B). The obvious hypothesis is that stress triggers additional condensation processes beyond translation initiation blockade alone. While we do not yet know which molecules are responsible for this additional stress-induced mRNP condensation, multiple RNA-binding proteins have been shown to autonomously sense heat shock and undergo condensation ^7,35,36,38^.

We then asked how pharmacologically blocking SG formation affects mRNP condensation. Treatment with cycloheximide (CHX) prior to stress prevents stress granule formation ^38,68,69^, which we confirm at 46°C heat-shock (Figure 7C). There is a clear contrast between inhibiting translation initiation (via depletion of eIF3b) and inhibiting translation elongation (via CHX): the former triggers SGs, while the latter prevents SGs.

However, CHX treatment sufficient to prevent SG formation reduces mRNP condensation only slightly (Figure 7D), consistent with persistence of TIICs. Are TIICs merely P-bodies (PBs), the other major cytosolic mRNP condensate known to be associated with translationally repressed mRNAs ^70^? Like SGs, PBs are also disrupted by CHX ^71,72^, while TIICs survive CHX treatment. Moreover, any stress which leads to translation initiation blockade should enhance TIIC formation, such that major PB proteins should accumulate in sedimentable assemblies. However, Lsm proteins, which are core PB proteins present at high concentration in PBs and which assemble into PBs in vitro ^73–75^, do not form sedimentable condensates during 46°C heat shock (Fig. S7), unlike TIICs. It remains possible that some subset of SG or PB-associated proteins form TIICs, consistent with the overlap between these two structures and our finding that TIIC accumulation potentiates SG formation. A recent study of acute glucose withdrawal also finds that translationally repressed mRNAs are largely found outside of P bodies ^16^.

We conclude that inhibiting SGs does not prevent mRNP condensation, and that TIICs are not SGs or PBs.

## Discussion

What is the physiological role of mRNP condensation in and outside of stress? Which mRNPs condense during stress, and why? What is the relationship between mRNP condensation, its functional causes and consequences, and stress granule formation?

We find that, across multiple stress conditions, preexisting mRNAs enter translationally silent condensates to a degree which depends on stress intensity. At the same time, stress-induced transcripts escape condensation and are robustly translated. These results echo early observations that stress granules exclude bulk nascent mRNA ^11,12^ and specific stress-induced heat shock protein transcripts ^13,14^. Our studies reveal that the timing of transcript production, rather than any particular transcript feature, is a primary determinant of escape from condensation; demonstrate the escape of dozens of stress-specific transcripts; and show that this escape from condensation permits selective translation. An important result from our study is that stress granules *per se* play little if any role in these processes.

### Small mRNP condensates are pervasive in the absence of stress or stress granules

Using a range of approaches, we discover pervasive mRNP condensation in cells without stress granule formation, and even in the absence of any discernible stress. Our results illuminate a level of molecular organization governed by translation initiation: initiation-blocked transcripts enter into structures we term translation-initiation-inhibited condensates (TIICs). TIICs can be generated containing specific mRNAs by blocking message-specific initiation, or at the transcriptome scale by blocking initiation; they do not require environmental stress for their formation; and they can form when stress granules are either absent or are pharmacologically blocked. This latter result mirrors the persistence of condensates of poly(A)-binding protein when stress granules are blocked ^38^. In short, TIICs are not stress granules.

In our experiments, we make no attempt to isolate stress granules *per se*. Given that a range of stress conditions—physiological stresses such as 42°C heat shock and 5% ethanol, and the less physiologically relevant but widely used 0.5% sodium azide—do not produce stress granules in our hands, but do produce considerable mRNP condensation, considerable biology would be overlooked by focusing only on SG-forming conditions. We show that mRNP condensation, and specifically TIIC formation, precedes and potentiates stress granule formation, and we confirm by single-molecule FISH that stress-induced transcripts escape from stress granules. Overall, our results support a model in which stress-associated inhibition of translation initiation causes formation of TIICs which, under intense stress, further assemble into stress granules by separate processes.

### mRNP condensation in cells is not primarily driven by ribosome-free RNA

Stress granules have long been thought to form after translation inhibition and ribosome runoff, exposing ribosome-free RNA which serves as a platform for new intermolecular interactions, whether directly between RNAs or mediated by RNA-binding proteins.^4,76–78^ This model has been strongly informed by results showing stronger recruitment of longer mRNAs and the ability of bound ribosomes to prevent SG recruitment.^79,80^ We find that the recruitment of initiation-blocked mRNPs into TIICs prior to, and independent from, SG formation proceeds quite differently. Length has little effect. Two abundant mRNAs, *HAC1* and *GCN4*, both with long stretches of ribosome-free mRNA, show divergent behavior: initiation-blocked *HAC1* mRNA condenses while uORF-regulated *GCN4* mRNA remains largely uncondensed. We reproduce these behaviors using synthetic mRNAs, isolating the critical role of blocked initiation rather than ribosome runoff for TIIC formation. Accordingly, locking ribosomes on mRNAs using CHX does not prevent TIIC formation.

In the context of stress granules, we can articulate two models for the effect of ribosomes on recruitment. In the first model, ribosomes prevent exposure of naked mRNA that is required for recruitment to SGs. In the second, ribosomes inhibit the processes that recruit mRNAs to SGs, and naked mRNA plays little or no role. We show that global translation initiation blockade and subsequent ribosome runoff from virtually all transcripts does not cause SG formation, falsifying the first model. Consistently, in mammalian cells, the presence of a single ribosome on an mRNA is sufficient to prevent recruitment even when the coding sequence is ribosome-free.^79^

Together, both for TIICs and for stress granules, ribosome-free mRNA appears to play a negligible causal role in the formation of these condensates.

### A protein-mediated cap-competition model coherently explains multiple mRNP condensate phenomena

Instead, our data consistently implicate 5’ end-mediated mRNA recruitment, although 3’ effects may play a role. We propose that competition by different processes for the mRNA 5’ cap can explain all of our observations (Figure 7E). Central to this model is the (presumably protein-mediated) recruitment of mRNAs to TIICs via binding to free 5’ cap. Such a model explains why mRNA length has little influence on TIIC formation, as well as why the presence of ribosomes in the mRNA body do not disperse TIICs. Any process that blocks access to direct cap binding would interfere with TIICs in this model. For most mRNAs, binding of the cytosolic eIF4F complex containing cap-binding protein eIF4E is transient, stabilized by mRNA activation and translation initiation.^81^ Consequently, most mRNAs will transiently have eIF4E-unprotected caps, making them substrates for condensation, and stable protection will be conferred by translation initiation. This latter fact would explain why we consistently observe tradeoffs between initiation and condensation, whether at steady state, for synthetic constructs, or when initiation is blocked by stress or by depletion of initiation factors.

### How do newly synthesized mRNAs escape condensates?

The cap-competition model also strongly hints at a mechanism, as yet undetermined, by which newly synthesized mRNAs may escape condensation. Our transcriptomic and reporter assays both show that transcripts transcribed during stress escape condensation regardless of sequence-encoded mRNA features or regulation by particular transcription factors. Consistent with our conclusion that timing is the key variable, an independent study of glucose withdrawal, another stress known to promote stress granule formation, also shows that expression timing, rather than sequence, determines whether mRNAs escape stress-induced translational repression.^16^

One possible explanation for the role of timing is that the force driving condensation in the cytosol weakens over the course of the stress, allowing subsequently exported transcripts to remain uncondensed. Alternatively, new transcripts may be marked in some way before or during nuclear export, blocking condensation while permitting translation initiation. Translation is not required for exclusion of new transcripts, because even when translation is inhibited by depletion of eIF3b, newly transcribed transcripts still escape. What might this condensation-inhibiting mark be? Possibilities include an mRNA modification such as methylation (or its stress-induced absence), changes in polyadenylation, or binding of a protein factor. Notably, most capped mRNAs are protected by a nuclear cap-binding complex after transcription and during export. This complex may be stabilized in the cytoplasm during stress instead of being exchanged during a pioneer round of translation, as suggested by work showing that nuclear cap-binding proteins can support active translation during stress.^82^ Such a complex would naturally prevent cap-dependent condensation, thus privileging newly synthesized mRNAs.

### What are the functions of mRNP condensation?

In light of our results, an accounting of the cellular function of mRNP condensation must contend with four facts: the presence of condensation in unstressed cells, the strong causal link to translation initiation inhibition, the weak dependence on mRNA length or sequence, and the exclusion of stress-induced messages. Exclusion of new messages and condensation of older messages also strongly favors an adaptive interpretation: stress-induced mRNP condensation helps cells rapidly redirect translational activity to transcripts most relevant to the cell’s current situation. This functional interpretation contrasts with previous work—heavily informed by the apparent length-dependence of condensation—questioning whether some RNA condensates are simply incidental products of translational inhibition during stress^19^ or, even more strongly, related to an “RNA entanglement catastrophe” resulting from overwhelming the RNA chaperoning capacity in the cell.^67,83^

We hypothesize that mRNP condensation provides cells with regulatory control over the translationally active transcriptome through a simple mechanism: preventing reinitiation of ribosomes on translationally stalled mRNAs by sequestering their 5′ ends in a condensate. Condensation, by sequestering transcripts away from competitive processes such as decapping or reinitiation,^84^ stores these mRNAs for short-term retrieval by dispersal factors including molecular chaperones. Blocking reinitiation is crucial for redirecting translational activity, and separable from another effect which is implied: protection of mRNAs from degradation.^26,85,86^ which would otherwise be another mechanism to prevent reinitiation. No part of this regulatory model requires formation of visible stress granules or other large membraneless organelles.

Beyond separating mRNP condensation from stress granule formation, a key advance reported here is to separate mRNP condensation from stress itself. How TIICs form, dissolve, influence regulation, and so on outside of stress must now become a focus.

## Data and code availability

All raw sequencing data generated for this project have been deposited in GEO under accession code GSE265963. All other data and code is deposited at https://github.com/drummondlab/RNACondensation2025/ (doi:10.5281/zenodo.15635227) or available upon request.

## Methods

### Cell growth and stress conditions

Unless otherwise noted, the BY4741 strain of *Saccharomyces cerevisiae* was used in experiments. All experiments were done with at least two biological replicates, starting from growth. Cells were grown at 30°C in synthetic complete dextrose media (SCD) for at least 12 hours to OD_600_ = 0.4 before being exposed to stress. Temperature stresses for sedimentation experiments were completed by centrifuging the culture and exposing the yeast pellet to either 42°C or 46°C water baths. Control cells were placed inside a 30°C incubator. Cycloheximide treated cells were pre-treated for 10 minutes with 100 μg/mL cycloheximide (Sigma #C7698-5G) before heat shock. Azide stresses were completed at either 0.5% w/v or 0.8% w/v for 30 min in SCD adjusted to pH 6.8 with NaOH. Azide was added from a 10% w/v sodium azide stock in water. Mock treatments were completed by adding pure water at the same volume to cultures. Ethanol stresses were completed by resuspending centrifuged cell pellets in SCD made with either 5%, 7.5%, 10%, or 15% ethanol for 15 min. Control cells were mock treated by resuspending in normal SCD. DTT treated cells were treated with 10 mM DTT for 15 minutes prior to harvesting. Temperature stresses for polysome sequencing and for tet-inducible reporter experiments were done by growing 250 mL of yeast in SCD overnight to OD_600_ = 0.4, collecting yeast via vacuum filtration onto a 0.45 μm filter (Cytiva 60206), putting the filter in 125 mL of pre-warmed media and incubating in a temperature controlled shaking water bath or incubator. After the indicated time, samples were harvested again via vacuum filtration and immediately scraped into liquid nitrogen.

Yeast transformations were performed either using a standard lithium acetate transformation or Zymo Frozen-EZ Yeast Transformation II Kit (Zymo #T2001) before plating on appropriate selection media ^87^. Clones were verified by colony PCR and Sanger sequencing.

### Generation of spike-in RNA

In-vitro transcribed (IVT) RNA or purified *Schizosaccharomyces pombe* total RNA was used as spike-ins where noted. The IVT RNA was produced by first amplifying a linear DNA fragment encoding NanoLuc using Q5 polymerase (NEB #M0494S), and purifying the DNA using an NEB clean and concentrate kit. The RNA was then made using a T7 Highscribe kit (NEB #E2040S), treated with DNase I (NEB #M0303L) and purified using an NEB clean and concentrate kit (NEB #T2030).

For the *S. pombe* RNA, fission yeast (FY527) was grown in YES media (5 g/L yeast extract, 30 g/L glucose, 225 mg/L adenine, histidine, leucine, uracil and lysine hydrochloride) at 32°C until OD_600_ = 0.5, harvested by centrifugation (3 minutes at 2500 g), resuspended in Trizol, and lysed by vortexing with 0.5 mm zirconia glass beads before extracting RNA using Zymo Direct-zol kits (Zymo #R2072).

### Fractionation-by-sedimentation sequencing (Sed-seq)

Biochemical fractionation was completed similarly to Wallace *et al*. ^38^, with the major exception that 20,000 g for 10 min was used rather than the original 100,000 g for 20 min. In short, 50 mL cultures of treated yeast were harvested by centrifugation at 3000 g for 5 minutes, then resuspended in 100 µL of soluble protein buffer (SPB: 20 mM HEPES, pH 7.4, 140 mM KCl, 2 mM EDTA, 0.1 mM TCEP, 1:200 protease inhibitor (Millipore #539136), 1:1000 SUPERase•In RNase Inhibitor (Invitrogen #AM2696), and flash frozen in liquid nitrogen as a pellet in a 2 mL Eppendorf Safe-Lock tube (Eppendorf #0030123620) with a 7 mm steel ball (Retsch #05.368.0035). The cells were then lysed using a Retsch MM400 for 5×90s at 30 Hz, chilling in liquid nitrogen between each shaking repeat. The lysed cells were resuspended in 600 µL of SPB, and 100 µL of total sample was transferred to 300 µL of Trizol LS (Invitrogen #10296010). For the S. pombe spike-in experiment, purified S. pombe total RNA was added to the lysate immediately after resuspension in SPB. The remainder was centrifuged for 30 seconds at 3000 g, and 300 µL of clarified lysate was transferred to a new 1.5 mL tube. This was then centrifuged for 10 minutes at 20,000 g. A 100 µL supernatant sample was transferred to 300 µL of Trizol LS, and 400 µL of SPB was added to the pellet as a wash. After another spin at 20,000 g for 10 minutes, the supernatant was removed and the pellet was resuspended by vortexing for 15 minutes in 300 µL of Trizol LS and 100 µL of water. If required, 1 ng of spike-in transcript was added to each sample at this step before RNA was isolated using Zymo Direct-Zol RNA extraction columns (Zymo #R2052), and RNA integrity was assessed by the appearance of two sharp rRNA bands on a 1% agarose gel and quantified using the absorbance at 260 nm.

### RNA quantification by RT-qPCR

Reverse transcription for qPCR was either performed using gene-specific reverse priming with the iScript™ Select cDNA Synthesis Kit (Bio-Rad #1708897) or using NEB LunaScript RT SuperMix kit (NEB #E3010L). In both cases, manufacturer protocols were followed using an input of 2.5 ng of RNA per µL of reaction. For gene-specific priming, the reverse primer was used at 5 µM. The IDT Primetime gene expression master mix (IDT #1055771) was used for quantitative PCR on a Bio-Rad CFX384 instrument with Taqman probes (1.5 µM for primers; 600 nM probe). For samples with spike-ins, abundances were calculated relative to the spike-in abundance using the ΔΔCq method.

### Polysome collection and analysis

Around 100 mg of frozen yeast that was collected by vacuum filtration, or following centrifugation at 3,000g for 1 minute, was transferred to a pre-chilled 2 ml Eppendorf “Safe-Lock” tube. Cells were lysed with a pre-chilled 7 mM stainless steel ball (Retsch #05.368.0035) by 5×90s×30Hz pulses in a Retsch MM100 mixer mill, chilling in liquid nitrogen (LN2) between pulses. Sample was resuspended in 10:1 (v/w) polysome lysis buffer (20 mM HEPES-KOH (pH 7.4), 100 mM KCl, 5 mM MgCl2, 200 μg/mL heparin (Sigma #H3149), 1% triton X-100, 0.5 mM TCEP (Goldbio #TCEP25), 100 μg/mL cycloheximide (Sigma #C7698-5G), 20 U/ml SUPERase•In (Invitrogen #AM2696), 1:200 Millipore protease inhibitor IV #539136). For the hairpin experiments in unstressed cells, samples were resuspended in polysome lysis buffer lacking heparin. For EDTA experiments, samples were resuspended in polysome lysis buffer lacking heparin and cycloheximide with 40mM EDTA to chelate Mg^2+^ and disrupt ribosomal complexes. The lysate was clarified by centrifugation at 3000 g for 30 s, and the clarified lysate was transferred to a new tube and aliquots were flash frozen in LN2.

A 10–50% continuous sucrose gradient in polysome gradient buffer (5 mM HEPES-KOH (pH 7.4), 140 mM KCl, 5 mM MgCl2, 100 μg/ml cycloheximide, 10 U/ml SUPERase•In, 0.5 mM TCEP) was prepared in SW 28.1 tubes (Seton #7042) using a Biocomp Gradient Master and allowed to cool to 4°C. Clarified lysate (200 µL) was loaded on top of the gradient, and gradients were spun in a SW28.1 rotor at 28,000 rpm for 3.5 hr at 4°C. Gradients were fractionated into 0.6mL fractions using a Biocomp Piston Gradient Fractionator with UV monitoring at 254 nm, and fractions were flash frozen in LN2. UV traces were normalized to the total signal starting with the 40S peak.

The samples were generated by pooling 50 µL of each fraction from the free fraction (before the monosome peak) and either separately pooling the fractions with 3+ ribosomes bound and the mono/di-some fractions (for the heat shock experiments), or by combining all ribosome-bound fractions together (azide and ethanol stresses). For the hairpin experiments in unstressed cells, samples were generated by pooling 75 µL from each pair of adjacent gradient fractions. The spike-in (50 ng of S. pombe total RNA) was then added to each pooled sample. RNA was purified via ethanol precipitation (final concentrations of 0.3 M sodium acetate pH 5.2, 0.3 µg/mL glycoblue (Invitrogen #AM9516), and 70% ethanol) at-20°C overnight followed by centrifugation at 4°C for 30 minutes at 21,000 g. The pellet was washed with 1 mL of 70% ethanol before being resuspended in water. The purified RNA was then treated with Dnase I (NEB) before purifying again using an NEB RNA clean and concentrate kit (NEB #T2030).

### Membrane flotation assay

Assay was performed with some modifications based on previous work ^64^. ∼100 mg of frozen yeast were transferred to a pre-chilled 2 ml Eppendorf “Safe-Lok” tube and lysed with a pre-chilled 7 mM stainless steel ball (Retsch #05.368.0035) by 5×90s×30Hz pulses in a Retsch MM100 mixer mill. Cells were chilled in liquid nitrogen (LN2) between pulses. Samples were resuspended in 10:1 (v/w) lysis buffer (20 mM HEPES-KOH (pH 7.4), 140 mM KCl, 5 mM MgCl2, 100 μg/ml cycloheximide, 10 U/ml SUPERase•In, 0.5 mM TCEP, 1% triton X-100). Lysis buffer was made lacking 1% triton X-100 when indicated. Lysate was clarified by centrifugation at 3,000g for 30 s and 250 µL of supernatant was mixed with 500 µL of 60% Optiprep iodixanol (Axis-shield). From this mixture, 600 µL were collected and dispensed at the bottom of SW 55 Ti tubes (Beckman Coulter #349622). Each tube was filled with 1.4 mL of 30% Optiprep with 100 µL of lysis buffer loaded on top. Samples were spun in SW 55 Ti rotor at 55,000 rpm for 2.5 hr at 4°C. Following centrifugation, gradients were manually fractioned starting from the top into 6 fractions of 350 µL. For each fraction, 50 µL was boiled in 2x Laemmeli buffer and 150 µL had a spike-in (50 ng of S. pombe total RNA) added prior to RNA purification via ethanol precipitation (final concentrations of 0.3 M sodium acetate pH 5.2, 0.3 µg/mL glycoblue (Invitrogen #AM9516), and 70% ethanol) at-20°C overnight followed by centrifugation at 4°C for 30 minutes at 21,000 g. The resulting pellet was resuspended in water and treated with DNase I (NEB) before being purified using an NEB RNA clean and concentrate kit (NEB #T2030).

### Sucrose cushion ribosome occupancy analysis

The ribosome occupancy (fraction of mRNA bound to ribosome) for the induction reporters was measured by spinning lysate through a sucrose cushion. Around 100 mg of frozen yeast was transferred to a pre-chilled 2 ml Eppendorf “Safe-Lok” tube. Cells were lysed with a pre-chilled 7 mM stainless steel ball (Retsch #05.368.0035) by 5 x 90s x 30Hz pulses in a Retsch MM100 mixer mill, chilling in liquid nitrogen (LN2) between pulses. Sample was resuspended in 10:1 (v/w) polysome lysis buffer (20 mM HEPES-KOH (pH 7.4), 100 mM KCl, 5 mM MgCl2, 200 μg/mL heparin (Sigma #H3149), 1% triton X-100, 0.5 mM TCEP (Goldbio #TCEP25), 100 μg/mL cycloheximide (Sigma #C7698-5G), 20 U/ml SUPERase•In (Invitrogen #AM2696), 1:200 Millipore protease inhibitor IV #539136). The lysate was clarified by centrifugation at 3000 g for 30 s, and 500 µL clarified lysate was transferred to a new tube.

At this point the sample was split into +/-EDTA samples. For the +EDTA samples, 6 µL of 0.5 M EDTA (pH 8 in water) was added to 150 µL of clarified lysate and incubated on ice for 10 minutes. Then 100 µL of both samples (+/- EDTA) was gently added on top of 900 µL of matching sucrose cushion (5 mM HEPES-KOH (pH 7.4), 140 mM KCl, 5 mM MgCl2, 100 μg/ml cycloheximide, 10 U/ml SUPERase•In, 0.5 mM TCEP, 20% sucrose w/v, +/- 20 mM EDTA) and centrifuged for 60 minutes at 100,000 g in a TLA55 rotor (Beckman-Coulter) at 4°C. The top 250 µL of supernatant was removed as the supernatant sample and 100 µL of this was mixed with 300 µL Trizol LS. The remaining supernatant was discarded before resuspending the pellet in 100 µL water + 300 µL Trizol LS (pellet is 10x relative to supernatant). To the pellet 1 ng of spike-in RNA was added, but only 0.1 ng was added to the supernatant.

RNA was purified from the supernatant and pellet samples using Zymo Direct-Zol kits, then the abundances of target RNAs were quantified via qPCR as above. Ribosome occupancies were calculated by calculating the percentage of each transcript in the pellet, after correcting for the pelleting observed in the presence of EDTA (this separates EDTA-sensitive polysomes in the pellet from EDTA-insensitive condensates).

### RNA sequencing

In general, DNase I treated RNA was prepared for sequencing using rRNA depletion (Illumina RiboZero (Illumina #MRZY1306) or Qiagen FastSelect (Qiagen #334215) followed by NEB NEBNext Ultra II (NEB #E7760) or Illumina TruSeq library preparation and Illumina platform sequencing. Specific methods for library preparation, sequencing and initial data analysis are described below and the method used for each sample is indicated in Table S4.

### Sequencing analysis

#### Genome references

Saccharomyces cerevisiae reference genome files (S288C_reference_genome_R64-3-1_20210421) were downloaded from the Saccharomyces Genome Database ^88^. Schizosaccharomyces pombe reference genome files were downloaded from PomBase^89^. When appropriate (see Table S4), the sequences of the NanoLuc spike-in or the mCherry and Clover reporters were included in the genome and transcriptome files for mapping.

#### Group A (see Table S4)

Sequencing libraries were prepared by the University of Chicago Genomics Facility from DNase I treated RNA using Illumina RiboZero (Illumina #MRZY1306) and Illumina TruSeq library prep kits. Single end 50 bp sequencing was performed on an Illumina HiSeq 4000 sequencer.

Sequencing reads were trimmed using TrimGalore (v0.6.10, https://github.com/FelixKrueger/TrimGalore) using default settings (e.g. trim_galore –gzip--fastqc_args’--outdir fastqc/’-j 4-o trimmed--basename FW32 EW_FW32_R1.fastq.gz). They were mapped using STAR v2.7.10b^90^ (e.g. STAR--outSAMtype BAM Unsorted--readFilesCommand gunzip-c--sjdbGTFfile saccharomyces_cerevisiae_R64-3-1_20210421_nofasta_geneid.gff--sjdbGTFtagExonParentTranscript Parent--sjdbGTFfeatureExon CDS--sjdbGTFtagExonParentGene gene_id--runThreadN 4--alignMatesGapMax 20000--limitBAMsortRAM 1445804817--genomeDir STAR_saccharomyces_cerevisiae_R64-3-1_20210421_allchrom –outFileNamePrefix mapped_reads/FW32/FW32_--readFilesIn trimmed/FW32_trimmed.fq.gz). To generate estimated counts and transcript per million (TPM) values, sequencing reads were mapped to the yeast transcriptome using kallisto v0.48.0^91^ (e.g. kallisto quant-i Scerevisiae_orf_coding_all_Scerevisiae_rna_coding.fasta.idx-o kallisto_quant/FW32--single-l 200-s 1--rf-stranded--bootstrap-samples=50-t 1 trimmed/FW32_trimmed.fq.gz).

#### Group B (see Table S4)

Sequencing libraries were prepared by from DNase I treated RNA using Qiagen FastSelect (Qiagen #334215), NEBNext Multiplex Oligos (UMI Adaptor RNA Set 1, NEB #E7335L) and NEBnext Ultra II Directional RNA library prep kits (NEB #E7760L). Paired end 200 bp sequencing with additional reads for dual 8/8 indices plus the 11nt UMI after the i7 index was performed on an Illumina NovaSeq 6000 at the University of Chicago Genomics Facility.

The unique molecular indices (UMIs) were extracted from fastq R2 using Umi-Tools v1.1.4^92^ and stored in fastq R1 and R3 (e.g. umi_tools extract--bc-pattern=XXXXXXXXNNNNNNNNNNN-I AD-JB-1S-HG02_S2_R2_001.fastq.gz--read2-in=AD-JB-1S-HG02_S2_R1_001.fastq.gz--read2-out=labeled_fastq/HG002/HG002_R1.umi.fastq. Sequencing reads were then trimmed using TrimGalore (v0.6.10, https://github.com/FelixKrueger/TrimGalore) using default settings (e.g. trim_galore--paired--gzip--fastqc_args’--outdir fastqc/’-j 4-o trimmed--basename HG002 labeled_fastq/HG002/HG002_R1.umi.fastq labeled_fastq/HG002/HG002_R3.umi.fastq). They were mapped using STAR v2.7.10b^90^ (e.g. STAR--outSAMtype BAM Unsorted--readFilesCommand gunzip-c--sjdbGTFfile spike_saccharomyces_cerevisiae_R64-3-1_20210421_geneid.gff3--sjdbGTFtagExonParentTranscript Parent--sjdbGTFfeatureExon CDS--sjdbGTFtagExonParentGene gene_id--runThreadN 4--alignMatesGapMax 20000--limitBAMsortRAM 1445804817--genomeDir STAR_spike_saccharomyces_cerevisiae_R64-3-1_20210421--outFileNamePrefix mapped_reads/HG002/HG002_--readFilesIn trimmed/HG002_val_1.fq.gz trimmed/HG002_val_2.fq.gz). Umi-Tools was then used again to deduplicate the reads (e.g. umi_tools dedup--stdin=mapped_reads/HG002/HG002_Aligned_Sorted.out.bam--chimeric-pairs=discard--unpaired-reads=discard--spliced-is-unique--paired-S mapped_reads/HG002/HG002_Aligned.sortedByCoord.dedup.out.bam). The reads were split again into fastq files using samtools v1.16.1^93^, and then estimated counts and TPMs were generated using kallisto v0.48.0^91^ (e.g. kallisto quant-i spike_Scerevisiae_orf_coding_all_Scerevisiae_rna_coding.fasta.idx-o kallisto_quant/HG002--rf-stranded--bootstrap-samples=50-t 1 mapped_reads/HG002/HG002_Aligned_dedup_R1.fastq.gz mapped_reads/HG002/HG002_Aligned_dedup_R3.fastq.gz).

#### Group C (see Table S4)

Sequencing libraries were prepared by the University of Chicago Genomics Facility from DNase I treated RNA using Qiagen FastSelect (Qiagen #334215) and Illumina Stranded mRNA Prep (Illumina #20020595) kits. Paired end 200 bp sequencing was performed on an Illumina NovaSeq 6000.

Sequencing reads were trimmed using TrimGalore (v0.6.10, https://github.com/FelixKrueger/TrimGalore) using default settings (e.g. trim_galore –paired--fastqc_args’--outdir fastqc/’-j 4-o trimmed--basename F02 AD-JB-F02_S44_R1_001.fastq.gz AD-JB-F02_S44_R2_001.fastq.gz). They were mapped using STAR v2.7.10b^90^ (e.g. STAR--outSAMtype BAM Unsorted--readFilesCommand gunzip-c--sjdbGTFfile spike_saccharomyces_cerevisiae_R64-3-1_20210421_geneid.gff3--sjdbGTFtagExonParentTranscript Parent--sjdbGTFfeatureExon CDS--sjdbGTFtagExonParentGene gene_id--runThreadN 4--alignMatesGapMax 20000--limitBAMsortRAM 1445804817--genomeDir STAR_spike_saccharomyces_cerevisiae_R64-3-1_20210421 –outFileNamePrefix mapped_reads/F02/F02_--readFilesIn trimmed/F02_val_1.fq.gz trimmed/F02_val_2.fq.gz). The estimated counts and TPMs were generated using kallisto v0.48.0^91^ (e.g. kallisto quant-i spike_Scerevisiae_orf_coding_all_Scerevisiae_rna_coding.fasta.idx-o kallisto_quant/F02--fr-stranded--bootstrap-samples=50-t 1 trimmed/F02_val_1.fq.gz trimmed/F02_val_2.fq.gz).

### Calculation of pSup

Public code for calculating pSup from sequencing data is available here: https://github.com/jabard89/sedseqquant. The statistical model used to estimate the proportion in supernatant (pSup) was based on that used in Wallace et al. (2015) ^33^. For each fractionated sample, the number of counts of mRNA within each fraction—total (T), supernatant (S), and pellet (P)—were extracted from RNA-sequencing data (see [“Sequencing Analysis” section above]). While mRNAs are expected to obey conservation of mass in the original fractionated lysate (𝑇_𝑖_ = 𝑆_𝑖_ + 𝑃_𝑖_ for mRNA species i), this assumption does not hold in the ratios of abundances directly inferred from the data. Instead, for a particular experiment, 𝑇_𝑖_ = α_𝑆_𝑆_𝑖_ + α_𝑃_𝑃_𝑖_ where we refer to the per-experiment constants α_𝑆_ and α_𝑃_ as mixing ratios which reflect differential processing and measurement of individual fractions. In order to estimate mixing ratios, and thus recover the original stoichiometry, we assume conservation of mass for each mRNA in the sample, and then estimate the mixing ratios under this constraint using a Bayesian model ^94^. We assume negative binomial noise for each count measurement, and log-normal underlying distribution of mRNA abundance. Specifically, we model counts as follows:

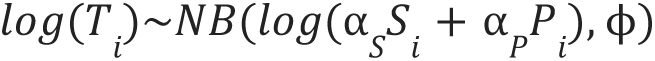

where

𝑇𝑖 = measured abundance of mRNA *i*,

𝑆𝑖 = measured abundance in supernatant of mRNA *i*,

𝑃𝑖 = measured abundance in pellet of mRNA *i*,

α_𝑆_ = mixing ratio of supernatant sample,

α_𝑃_ = mixing ratio of pellet sample

With the following priors:

α_𝑆_ ∼Γ(1, 1)

α_𝑃_ ∼Γ(1, 1)

σ∼𝐶𝑎𝑢𝑐ℎ𝑦(0, 3)

We implemented the model above in R using the probabilistic programming language STAN, accessed using the rstan package ^95,96^ and used all mRNA with 𝑐𝑜𝑢𝑛𝑡𝑠 > 20 to estimate mixing ratios for each sample. These mixing ratios were then used to calculate the pSup for mRNA *i*:

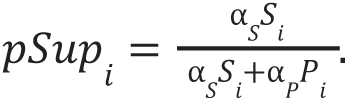

### Differential sedimentation and escape scores

To calculate the differential sedimentation (ΔSed) and escape (eSed) scores, which capture a stress-dependent difference from the treatment, we first calculate a windowed mean over transcript length of log-odds pSup in the mock (untreated) condition. Each window is 0.02 of the full range of transcript lengths on a log scale.

For a transcript with length L, we take the mean of the log-odds pSup values for all transcripts within a window centered on log L; these means μ(L,T) are calculated for all values of L in the transcriptome. We then compute the standard deviation σ of all transcript pSup values from the windowed mean for the corresponding length.

Given the resulting quantities:

lopSup(x,T) = log-odds pSup [log p/(1−p) if pSup = p] of gene x after treatment T μ(L,T) = mean lopSup in window around length L after treatment T,

σ(L,T) = standard deviation lopSup in window around length L after treatment T, σ(T) = standard deviation of lopSup(x,T = control) − μ(L,control) over all genes x

the differential sedimentation ΔSed for gene x, with transcript length L, after treatment T is

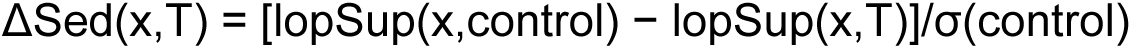

and the escape from sedimentation eSed for gene x after treatment T is

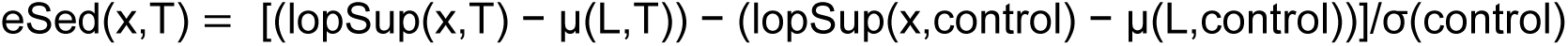

Intuitively, ΔSed captures changes in sedimentation due to treatment in units of σ(control). ΔSed = 0 for the control condition. Escape score eSed captures the treatment-induced difference in sedimentation relative to transcripts of the same length; eSed > 0 indicates less sedimentation than the average (escape), and eSed < 0 indicates more sedimentation than the average.

### Other bioinformatic analyses

#### Transcript features

Transcript features were extracted from Saccharomyces Genome Database (Cherry et al. 2012). Targets of HSF1 and MSN2/4 were based off those reported in Pincus et al. 2018^44^ and Solis et al. 2016^45^. Transcript UTR lengths were taken as the median value reported by long read transcript sequencing in Pelechano et al. 2013^97^, or, when no data was reported, the median UTR length in yeast was used as the default. Pombe transcript lengths, including the lengths of the UTRs, was taken from PomBase ^89^.

#### Transcript abundance

The transcript abundance is reported as the geometric mean of the TPM value for two biological replicates, estimated by kallisto analysis of the Total fraction for each sample. Changes in transcript abundance were calculated using DeSeq2^98^.

#### sedScore calculation

In order to calculate sedScores, the pSup for each transcript was converted to a log-odds scale, and transcripts were arranged by their length (including UTRs), and then binned into groups of 100. For each transcript in the bin, the standard deviation from the mean within the bin was used to calculate a Z-score. Individual Z-scores from two biological replicates were calculated and then averaged together for the final reported sedScore.

#### Ribosome occupancy

Because Polysome-seq data was collected with spike-in values for each fraction (Total, Free, Mono/Poly), it is possible to calculate the absolute ribosome occupancy (% of a transcript which is bound to at least one ribosome) for each transcript. This value is calculated by normalizing transcript abundance for each fraction (TPMs output by kallisto) to the median abundance of the spike-in transcripts. All S. pombe spike-in transcripts with more than 100 estimated counts were used to calculate the spike-in abundance. The ribosome occupancy is then calculated as abundance_bound_/(abundance_bound_ + abundance_free_).

#### Ribosome association

In stressed samples, it is possible that condensed RNA pellets to the bottom of the sucrose gradient, making it difficult to calculate the absolute ribosome occupancy. Thus, for stressed samples, we calculate a “ribosome association” score which is TPM_rib. bound_/TPM ^99^. This metric is similar to “translation efficiency” scores calculated for ribosome profiling studies^61^. The change in ribosome association upon stress was calculated using DeSeq2 ^98^, similar to reported methods for calculating changes in translation efficiency using DeSeq2 ^100^.

#### RNA structure analysis

The sequence for the 5′ UTR + the first 20 nucleotides of the CDS was extracted using the 5′ UTR lengths described above from Pelechano et al. 2013 ^97^. The folding energy for each UTR was then calculated using RNAFold from the ViennaRNA package ^101^. Because the folding energy correlates directly with length, a normalized structure score was calculated for each transcript by dividing the calculated folding free energy by the length of the UTR.

#### Inducible reporter genes

Reporters for pulsed induction were generated by Gibson assembly of gene fragments with a TET-inducible promoter designed for tight control of induction levels ^51^. Assembly pieces were derived either from gene fragments ordered from IDT or Twist Biosciences or from PCR amplification of other plasmids. Fragments were assembled into backbones generated by golden gate cloning using protocols and plasmids from the Yeast Toolkit ^102^, and the plasmids were sequenced by overlapped Sanger sequencing. Plasmids were linearized with NotI prior to transformation.

The PMU1 reporter contains the 5′ UTR and 3′ UTR of the native PMU1 gene and the CDS is a fusion of the PMU1 CDS with nanoluciferase-PEST^103^. The HSP26 reporter contains the 5′ UTR and 3′ UTR of the native HSP26 gene, but the CDS is a fusion of the TPI1 CDS and nanoluciferase-PEST. The TPI1 fusion was used to avoid potential artifacts caused by a large pre-induction of HSP26 molecular chaperone and because TPI1 is well translated during stress and of a similar length (645 nt for HSP26 vs 745 nt for TPI1). Reporters were integrated at the HO locus using hygromycin selection in a strain of yeast containing a C-terminal auxin tag on Sui2, along with the inducible TIR1 ligase at the LEU locus, and the TetR protein at the his locus (see Table S1 for full genotype).

For induction of reporters concurrently with stress, 1 µM anhydrotetracycline (aTC, Cayman #CAYM-10009542-500) was added from a 10 mM stock prepared in DMSO at the beginning of the stress. For pre-induced samples, 0.1 µM aTC was added to yeast in SCD at OD_600_ = 0.2 and samples were incubated at 30°C for 45 minutes. Samples were then either washed 3x with SCD via centrifugation, or 1x via vacuum filtration before resuspending in prewarmed SCD. Stress was then initiated 30 minutes after washing had begun to ensure complete shutoff of reporter transcription. Samples were then fractionated as described above either using the Sed-seq protocol to calculate pSup or the sucrose cushion fractionation to calculate ribosome occupancy.

#### Engineering solubility reporters

Solubility reporters were engineered using the Yeast Toolkit [Lee et al., 2015] (see Table S1 and S2). Variable 5′UTRs were engineered depending on the construct and genetically integrated in front of two copies of Clover, all driven by the constitutive TPI1 promoter and with the TPI1 3′ UTR. Each reporter construct also had a copy of mCherry with a TPI1 promoter, 5′UTR and 3′UTR. This construct was inserted into the Leu2 locus with leucine selection.

Steady state protein levels were measured using flow cytometry by normalizing the Clover signal to the mCherry signal in each cell. Data was analyzed with a custom script using FlowCytometryTools in python and then exported and plotted in R. The Sed-seq protocol was used to measure the condensation behavior of each strain. Steady state mRNA levels were extracted from the Total sample of the Sed-seq experiment and translation efficiency was calculated as the steady state protein level divided by steady state RNA level.

### Auxin-mediated depletion strains

Auxin induced degron depletions were adapted from the approach in Mendoza-Ochoa et al. [2019]. In short, the endogenous protein of interest was genetically engineered to contain the degron tag in a strain of yeast in which a β-estradiol inducible TIR1 ligase had been genetically integrated at the LEU locus. Some of the strains contained the original Oryza sativa TIR1 (OsTIR1), while others used a variant engineered for more specificity OsTIR1(F74G) ^54^ as indicated in Table S1. The auxin-FLAG degrons were installed at either the 5‘ or 3‘ end of genes using CRISPR plasmids from the yeast toolkit. A PCR-generated DNA template was co-transformed with a Cas9 and gRNA containing URA3 selectable plasmid as previously described ^102,104^. The CRISPR integrations were verified by PCR and Sanger sequencing and the URA3 plasmid was removed by selecting for colonies which did not grow on URA plates.

For depletion experiments, yeast were grown at 30°C in YPD to OD_600_ = 0.1. To induce TIR1 ligase, 5 µM β-estradiol (10 mM stock in DMSO) or an equivalent volume of DMSO (for mock treatment) was added to each culture and they were incubated for 75 minutes. To induce degradation, either 100 µM of Indole-3-acetic acid sodium salt (Sigma #I5148, 250 mM stock in DMSO) or 5 µM of 5-Ph-IAA (Medchemexpress #HY-134653, 5 mM stock in DMSO) was added. After 2 hours of auxin exposure, cells were temperature treated and then harvested and fractionated as normal.

### Radiolabeling quantification of translation

Yeast cells were cultured overnight in YPD until they reached an OD_600_ = 0.1. Auxin-inducible yeast strains were then treated with beta-estradiol and auxin, as detailed above, then translation was measured following a published protocol^105^. After a 1.5-hour depletion period, 1 mL of sample was transferred to 1.5mL tubes, then 1 µCi/mL of mixed 35S-L-methionine and 35S-L-cysteine media were added to each sample (Perkin-Elmer EasyTag #NEG772002MC). Samples were incubated for 30 minutes at 30°C with shaking (15 minutes for heat shocks), then cells were treated with 200 µL of 50% trichloroacetic acid (TCA), chilled on ice for 10 minutes, heated at 70°C for 20 minutes, and cooled again for 10 minutes. The samples were subsequently collected on glass microfiber filters (Sigma #WHA1823025) loaded onto a vacuum manifold (Millipore #XX2702550), washed with 3x 5 mL 5% TCA and 2x 5mL 95% ethanol, and air-dried for at least 12 hours at room temperature. Filters were then immersed in scintillation fluid (Perkin Elmer #6013179), and radioactivity levels were quantified in “counts per minute” through liquid scintillation counting on a Tri-Carb machine.

### Western blotting

Western blots were performed as described in a published protocol ^106^. For each sample, 1mL of yeast culture was spun down at 2500 g for 2 minutes, and the pellet was resuspended in 50 µL of 100 mM NaOH. The samples were incubated for 5 minutes at RT, spun at 20,000g for 1min, and resuspended in 50 µL of 1x Laemmli buffer (Bio-rad #1610737) with 5% β-mercaptoethanol. Samples were then boiled for 3 minutes, clarified at 20,000 g for 2 minutes and 15 µL was loaded onto a 4-20% tris-glycine SDS-PAGE gel (Biorad #5671094). Proteins were then transferred to nitrocellulose (Sigma #10600001) using a wet transfer apparatus (Bio-rad #1704070). The membrane was blocked for 1 hour with 5% milk in TBST buffer, then incubated rocking overnight at 4°C with 1:3000 dilution of anti-FLAG antibody (Sigma #F1804) and 1:10,000 dilution of anti-PGK1 antibody (Invitrogen #459250) in 5% milk solution. Westerns were visualized using 1:20,000 dilutions of fluorophore conjugated secondaries (Licor #926-32212 and #925-68073) and visualized on a Licor Odyssey CLx. Band intensities were quantified in ImageJ and normalized to PGK1 signal. For Dpm1, primary incubation was done at 1:1000 for 48 hours.

### Fluorescence microscopy and stress granule quantification

Standard confocal microscopy was completed as in Wallace et al. [2015], generally using Pab1-Clover as the SG marker unless otherwise noted. Cells were grown to log-phase as previously described. 1mL of cells were transferred to 1.5mL Eppendorf tubes. For heat stress, cells were shocked in a heat block, spun down in a microfuge, and 950 uL of supernatant were removed. For azide stress, 10% (w/v) azide or water was added directly to the 1mL of cells as indicated. For ethanol stress, cells were spun down in microfuge and resuspended in media with appropriate amounts of ethanol. 1.5 uL of treated cells were then placed on a glass slide and imaged immediately. For AID treatment, cells were treated as described above, and were imaged immediately after a 2 hour exposure to Auxin. For cycloheximide treatment, cells were exposed to 100 ug/mL of cycloheximide for 10 minutes, stressed for 10 minutes, and then imaged immediately. Cells were imaged on an Olympus DSU spinning disc confocal microscope using a 100x 1.45 TIFM oil objective (PlanApo) and the FITC filter cube for the Clover fluorophore in Z-stacks. Representative images are maximum projections of the collected z-stacks. Maximum projection images of the cells were used to quantify the number of stress granules per cell using CellProfiler.

### Single-molecule fluorescence in situ hybridization (smFISH)

Custom Stellaris® RNA FISH Probes were designed against SSB1, SSA4, HSP104, and ADD66 by utilizing the Stellaris® RNA FISH Probe Designer (Biosearch Technologies, Inc., Petaluma, CA) available online at www.biosearchtech.com/stellarisdesigner (Table S3). Each Stellaris FISH Probe set was labeled with Quasar670 (Biosearch Technologies, Inc.). smFISH was done as previously described ^107,108^. Yeast cultures were grown to an OD of 0.3-0.4 in SCD, spun down at 3k g for 3 min. Cells were then suspended into 4mL of culture and Oregon Green HaloTag reagent (Promega #G2801) was added to a final concentration of 2uM. Cells were then resuspended and split into final cultures of 25 mL. Cells were then spun again at 3000g for 3min, and 23mL were removed, such that 2mL of media remained. Cells were then stressed as stated before. 19.85mL of pre-warmed media was then added to each falcon tube, and 3.15 mL of 4% paraformaldehyde (Electron Microscopy Services #15714) was immediately added. Cells were incubated at room temperature for 45 min at room temperature, gently rocking. Cells were spun down at 4°C and washed with ice-cold buffer B. Cells were resuspended into 1mL of Buffer B (1.2M sorbitol, 100mM KHPO4, pH = 7.5) then transferred to a 12-well plate. Cells were additionally crosslinked in a Spectrolinker UV Crosslinker at a wavelength of 254nm by exposure to 100 mJ/cm^2 twice with 1 min break in between ^109^. Cells were pelleted for 3min at 2000rpm and then resuspended into spheroplast buffer (1.2 M sorbitol, 100 mM KHPO4, pH = 7.5, 20mM ribonucleoside-vanadyl complex (NEB # S1402S), 20mM B-mercaptoethanol). 25U/OD of lyticase (Sigma #L2524-10KU) were added to each sample. Cell digestion was performed at 30°C and was monitored using a benchtop phase contrast microscope, such that cells were about 50%-70% digested. Digestion was stopped by spinning cells at 4°C for 3min at 2000 rpm and two washes twice in ice cold buffer B and resuspended in 1mL Buffer B. 250 uL of cells were placed onto a poly-L lysine coated coverslip and incubated at 4C for 1hr. Cells were washed with 2mL of Buffer B and then stored in ice-cold 70% ethanol for at least 3 hours. Coverslips were rehydrated in 2xSSC and then washed twice in pre-hybridization buffer (2x SSC + 5% formamide (Sigma #344206-100ML-M)) for 5 minutes each. A mixture of 0.125uL of 25uM smFISH probes, and 2.5uL of 10mg/ml yeast tRNA (Thermo #AM7119) and 2.5uL of 10mg/mL salmon sperm DNA was dehydrated in a Speedvac at 45°C. The dried pellet was rehydrated was resuspended in 25 μl hybridization mix (10% formamide, 2×SSC, 1mg/mL BSA, 10 mM Ribonucleoside–vanadyl complex (Thermo #15632011) and 5 mM NaHPO4, pH 7.5) and boiled at 95 °C for 2 min. 18uL of resuspended probes were spotted onto a piece of Parafilm and coverslips were placed cell-side down into hybridization mixture. Hybridization occurred at 37°C for 3 hours. Coverslips were then washed at 37°C for 15min in 2x SSC + 5% formamide, then in 2x SSC buffer, then 1xSSC buffer. They were then submerged in 100% Ethanol, dried, and then mounted into ProLong Gold antifade with DAPI (Thermo P36941).

### smFISH image acquisition and analysis

smFISH images were taken on a Nikon TiE microscope with a CFI HP TIRF objective (100x, NA 1.49, Nikon), and an EMCCD (Andor, iXon Ultra 888). Nikon TiE epifluorescent microscope. Samples were excited using the 647nm laser (Cobolt MLD) (∼15-20 mW for 200-300ms), poly-A FISH was imaged using the 561nm laser (Coherent Obis) (∼15-20 mW for 200-300ms), and Pab1-Halotag signal was imaged with a 488nm laser (Cobolt MLD) (∼10-15 mW for 200-300 ms), and DAPI (CL2000, Crystal Laser) (∼5-10 mW for 100 ms). Imaging of the nucleus was done using the 405nm laser and DIC images were taken as well. Z-stacks of 21 planes, 2uM thick were obtained. Images were analyzed using FISH-quant ^110^. Briefly, RNA spots were identified using big fish^110^. For the smFISH colocalization analysis, RNA spot intensities were normalized by dividing by the mean intensity of each cell. For each RNA spot, the mean Pab1 intensity in a 3×3 pixel square around the centroid was calculated. The Pab1 intensity was then measured for 100 random locations in the cell in 3×3 pixel locations. Finally, a distribution was calculated for both the random Pab1 signal and the Pab1 signal that corresponds to a RNA spot. The Z-score of the mean intensity of the Pab1 signal in a RNA spot compared to the Pab1 signal in a random spot was compared, and this is termed the ‘colocalization score’. Each Z-score is calculated independently for each cell to account for different background intensities, and the average shown is for every cell.

### Fitting of mRNA and mRNP condensation

The underlying biophysical model for pSup in the absence of condensation is 𝑝𝑆𝑢𝑝(𝑔) = 1 − β𝐿*_𝑔_^χ^* for a mRNA transcript encoded by gene 𝑔, of length 𝐿𝑔. In conditions where there is mRNA condensation, governed by parameter µ per-transcript and ν per-nucleotide, the model is: 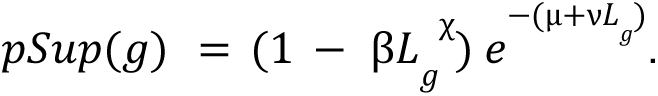 These models were fitted to sedimentation on the log-odds(pSup) scale, i.e. approximating the log-odds sedScore as normally distributed. Non-linear least squares fits were performed using the nls function in R. See supplemental text for details.

## Statistical analyses

Unless otherwise stated, all experiments were performed as at least two biological replicates, and the mean or geometric mean value (for log-distributed transcript abundance data) was calculated from the replicates. Unless otherwise noted, all correlation values are reported as Spearman’s rank correlation coefficient and significance tests comparing groups of data points were performed using a Wilcoxon rank-sum test, with a Bonferroni correction when multiple groups were being compared (*P < 0.05, **P < 0.01, ***P < 0.001.’N.S.’ denotes not significant (P ≥ 0.05).

## Acknowledgments

We thank Chris Katanski for help with experiments and extensive discussions. We thank Tineke Lenstra and Daniel Larson for discussions and help with smFISH, and Jean Beggs for providing the inducible TIR1 ligase system and for support and discussions on all things RNA. We thank Asif Ali for help making pJB773. FRP2371 was a gift from Joerg Stelling (Addgene plasmid #127577). pV1382 was a gift of Guy Bushkin and Gerald Fink. We are grateful to David Pincus, Rasi Subramaniam, and the entire Drummond group for numerous vital discussions and for helpful comments on the manuscript.

We thank Ross Buchan for sending us the Pbp1-GFP strain and helpful imaging suggestions. We thank Noah Mitchell and Kyle Lin for discussions of imaging quantification. We acknowledge the University of Chicago Integrated Light Microscopy Core (RRID: SCR_019197) and valuable assistance from the Biophysics Facility and the Genomics Facility at the University of Chicago.

We acknowledge funding from NIH (award R35 GM144278 to D.A.D., award F30 ES032665 to H.G., award F31 ES033554 to C.J.W.H., awards R01 GM055694 and R35 GM14833 to T.R.S., award R35 GM136296 to R.H.S; and award K99 GM148788 to W.L.), from European Union’s Horizon 2020 (Marie Skłodowska-Curie grant agreement no. 661179 to E.W.J.W.), Wellcome (208779/Z/17/Z to E.W.J.W.), and from the Helen Hay Whitney Foundation (award to J.A.M.B.). The content is solely the responsibility of the authors and does not necessarily represent the official views of the NIH.

