## Supplementary Information for "Transcriptome-wide mRNP condensation precedes stress granule formation and excludes new mRNAs"

#### Supplementary Text

##### A model of free mRNP sedimentation

Let $L$ be the length of an mRNA under consideration. We seek the functional form of $\mathrm{pSup}(L)$ , the proportion in the supernatant after centrifugation, which will depend on many factors. Several of these are experimental and we assume they do not change across samples or mRNAs, such as the spin speed $\omega$, the sample height $h$, the spin time $s$, and the viscosity $\eta$. Terminal velocity for a particle with mass $m$ and hydration radius $r_0$ is:

$$v_t(m, r_0) = \frac{m\omega^2 r}{6\pi\eta r_0} = \left(\frac{m}{r_0}\right)\left(\frac{\omega^2 r}{6\pi\eta}\right)$$

We assume that both mass $m$ and hydration radius $r_0$ scale consistently with length $L$ on average. In the case of mass, we assume proportional scaling: a constant (average) molecular weight per unit length deriving from nucleotides and bound proteins. In the case of hydration radius, our scaling assumption is consistent with standard approaches (Yoffe et al. 2008), for the radius of gyration $R_g$ scales as $L^\chi$ with $\chi = 0.34$ for naked ssRNA due to secondary structure. The hydration radius $r_0$ will be proportional to $R_g$ assuming a constant density. The scaling relationship will also depend on the geometry of the vessel.

Assuming a constant mass per nucleotide and protein binding per unit length, the mass of an mRNP scales linearly with $L$. Therefore, the leading term above will tend to scale with $L$:

$$\frac{m}{r_0} = (c_1 L)^\chi$$

with proportionality constant $c_1$ taking care of the specific conversion factors (e.g. the molecular weight per nucleotide) and with $\chi$ summarizing the scaling.

Then the proportion of the particle species remaining in the supernatant is the proportion of the tube which is more than $v_t s$ from the bottom. That is: $v_t s$ from the bottom. That is:

$$\mathrm{pSup}(m, r_0, h, s) = \begin{cases} 0 & v_t(m, r_0)s \geq h \\ 1 - \frac{v_t(m, r_0)s}{h} & 0 \leq v_t(m, r_0)s < h \end{cases}$$

We can then summarize the dependencies with two constants, as follows.

$$\mathrm{pSup}(L) = 1 - \frac{v_t(m, r_0)s}{h}$$

$$= 1 - \left(\frac{m}{r_0}\right)\left(\frac{\omega^2 r}{6\pi\eta}\right)$$

$$= 1 - c_2(c_1 L)^{\chi}$$

$$= 1 - \beta L^{\chi}$$

That is, two parameters should be sufficient to describe the average behavior of free (unclustered) mRNAs of length $L$. The model fits the average data well; see Figure 1E.

**A model of treatment-induced mRNP condensation**

To model stress-induced interactions and their effect on $\mathrm{pSup}$, we consider two kinds of interactions, both assumed to be Poisson-distributed with rates which vary across conditions and potentially across transcripts.

The first interaction type is between nucleotides, including direct RNA-RNA interactions and interactions mediated by proteins but dependent on nucleotide content; these are expected to be length-dependent. Consequently, we model them by a per-nucleotide interaction rate $\nu$, such that the probability that an mRNP has $k$ nucleotide-dependent interactions is

$$\Pr(k) = (\nu L)^k e^{-\nu L}/k!.$$

The second interaction type is between molecules and is independent of sequence length, including interactions mediated by the 5' cap or the 3' end. We model these as a per-molecule interaction rate $\mu$ such that the probability that an mRNP has $m$ such interactions is

$$\Pr(m) = \mu^m e^{-\mu}/m!.$$

These rates permit us to model the probability that an mRNP of a given length $L$ engages in interactions with other mRNPs, which may have varying lengths/sizes and may themselves also engage in additional interactions, making prediction of sedimentation highly complex. To proceed analytically, we make a simplifying assumption: that under the conditions we study, most mRNP condensates are sufficiently large to sediment completely (pSup = 0). Under this assumption, the observed proportion in the supernatant for an mRNA of length $L$ is given by the free-mRNP pSup multiplied by the probability that the mRNP is free. The latter probability is equal to the probability of $n = 0$ per-nucleotide interactions and $m = 0$ per-molecule interactions. Combining the results into a single expression, we have

$$\mathrm{pSup}(L) = (1 - \beta L^{\chi})\, e^{-(\mu + \nu L)}.$$

The mean behavior of transcripts in any particular experiment, in this model, can be derived from only four fitted parameters $\beta, \chi, \mu, \nu$ and the lengths of all transcripts. Fits of this model are shown in Fig. 1E, where the no-stress sample is modeled with no clustering ($\mu = 0$, $\nu = 0$ ) and stress samples are modeled with clustering (solid lines) and also in comparison to curves where the per-molecule interaction rate is zero ($\mu = 0$; dashed lines). Fitted values are shown in Figure S1E.

For given values of the two stress-induced-condensation-related parameters $\mu$ and $\nu$, the transcript length at which the rate of interactions is equal is given by $L\nu = \mu$ or

$$L_{\mathrm{equiv}} = \frac{\mu}{\nu}.$$

We can then ask what proportion of transcripts are at least this long, $L \geq L_{\mathrm{equiv}}$, such that the stress-induced condensation per-nucleotide (length-dependent) contributes more to condensation than per-molecule (length-independent) interactions. These results are tabulated in Figure S1E.

Models with and without a $\mu$ parameter are formally nested, and fits can be compared with an *F* statistic. For both 42°C and 46°C, the model without $\mu$ (all interactions are sequence-mediated) can be sharply rejected (*F* = 4134.66, *F* = 21,507.21, $P < 10^{-6}$ in both cases).

**Bounds on condensate sizes**

How can we put bounds on the average size of condensates, given an observed value of pSup? Any pSup value can be realized in a range of ways, bounded by two extremes. At one extreme, the molecular population is homogenous (all molecules of a particular type are in condensates of the same size), and all clusters have the same probability of remaining in the supernatant, which is pSup. At the other extreme, the molecular population is heterogeneous, with each species of condensate having some proportion in the supernatant $pSup_i$ and proportion of the molecular population $q_i$. Order the indices *i* such that the lightest species (monomers) has $i=1$ and all condensates are ordered by their size (and hence in the descending order of their pSup) up to *n*. First, if *n*=2, it is clear that the observed $pSup \geq pSup_2$, simply because the average of two numbers is greater than or equal to the smaller of the two numbers. Second, if we define the average of the condensates' pSups as $pSup_{cond}$, we have

$$pSup_{cond} = \frac{1}{1-q_1}\sum_{i>1}^{n} q_i\, pSup_i$$

So that

$$pSup = q_1\, pSup_1 + (1-q_1)pSup_{cond} \geq q_1\, pSup_{cond} + (1-q_1)pSup_{cond} = pSup_{cond}$$

which establishes $pSup \geq pSup_{cond}$ such that in general, pSup is an upper bound on the average condensate size.

### Supplementary Figures

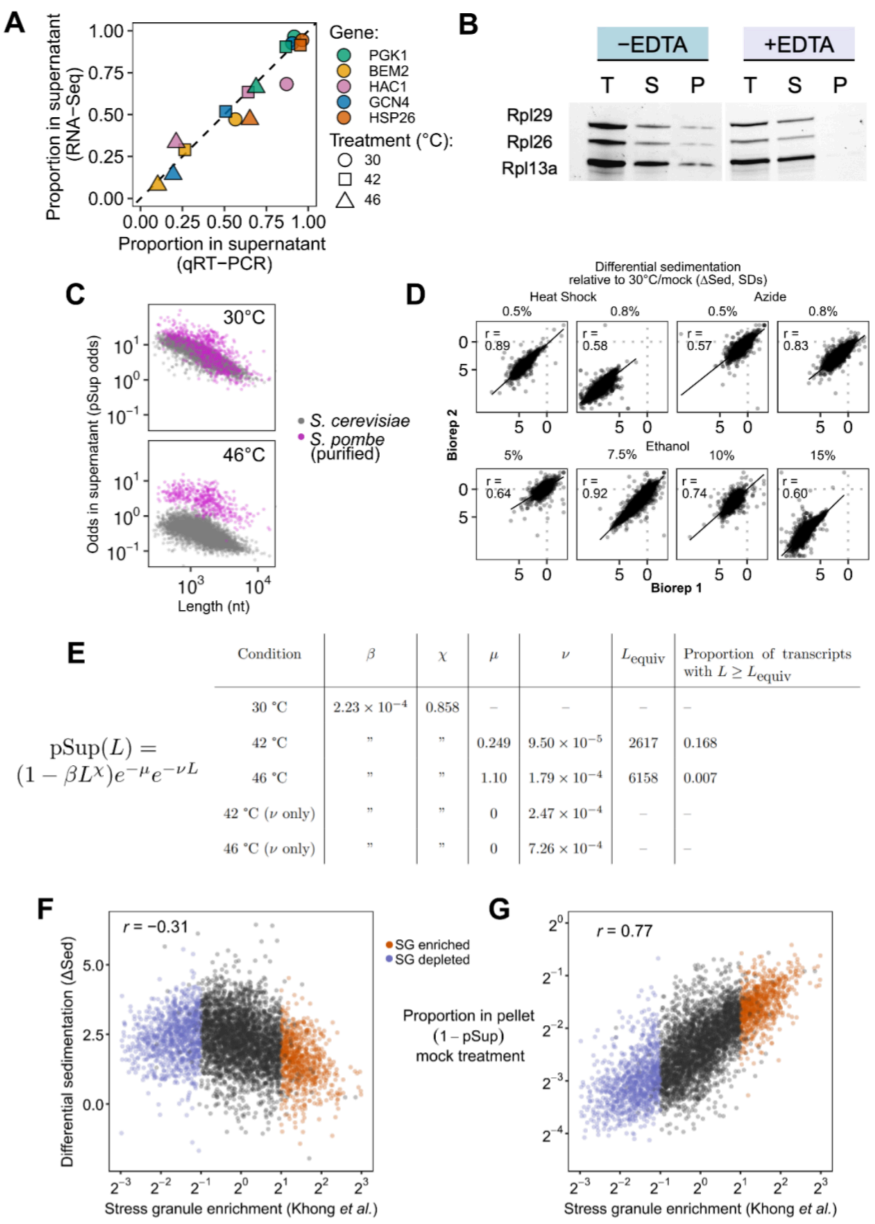

$$\mathrm{pSup}(L) = (1 - \beta L^{\chi})e^{-\mu}e^{-\nu L}$$

| Condition | $\beta$ | $\chi$ | $\mu$ | $\nu$ | $L_{\mathrm{equiv}}$ | Proportion of transcripts with $L \geq L_{\mathrm{equiv}}$ |
|---|---|---|---|---|---|---|
| 30 °C | $2.23 \times 10^{-4}$ | 0.858 | – | – | – | – |
| 42 °C | " | " | 0.249 | $9.50 \times 10^{-5}$ | 2617 | 0.168 |
| 46 °C | " | " | 1.10 | $1.79 \times 10^{-4}$ | 6158 | 0.007 |
| 42 °C ($\nu$ only) | " | " | 0 | $2.47 \times 10^{-4}$ | – | – |
| 46 °C ($\nu$ only) | " | " | 0 | $7.26 \times 10^{-4}$ | – | – |

**Figure S1**: **Sed-seq captures previously unreported mRNA condensation largely driven by length-independent mechanisms.** (A) The measurements of mRNA proportion in supernatant (pSup) from RNA sequencing match pSups calculated using spike-in-normalized abundances quantified by RT-qPCR. (B) EDTA prevents sedimentation of polysomes, as shown by western blot against three ribosomal proteins. See also Fig. 5G. (C) Purified mRNA (*S. pombe* total RNA) spiked into lysate from unstressed yeast cells recapitulates length-dependent sedimentation of free mRNA, and remains largely free even when spiked into lysate from severely stressed yeast cells. (D) Across stresses, differential sedimentation (ΔSed) of biological replicates correlate well *(r*, Pearson correlation coefficient). (E) Fitted parameters for the four-parameter model shown; see Supplementary Text. (F) The reported enrichment of transcripts in azide-induced stress granules (Khong *et al.* 2017) and our measured differential sedimentation transcript sedimentation during azide stress show mild negative correlation. (G) Most of the variation in stress-granule depletion in Khong *et al.* 2017 can be explained by sedimentation of free mRNPs due to their length, and thus can be closely reproduced using only Sed-seq data from unstressed (mock-treated) cells.

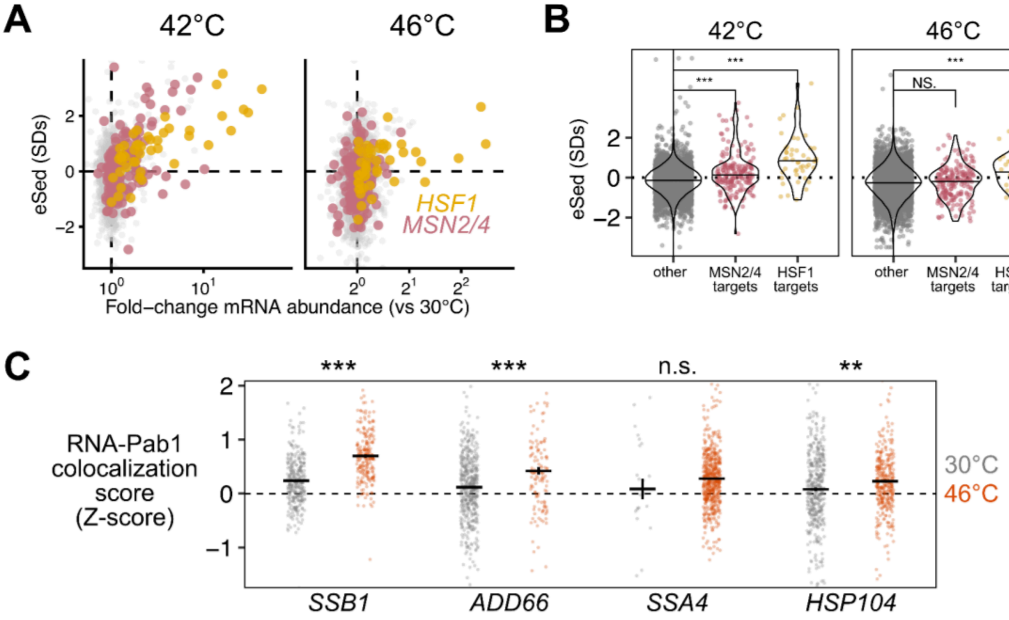

**Figure S2**: (A) Induced Hsf1 targets and Msn2/4 targets relatively escape condensation. (B) Hsf1-regulated transcripts escape condensation at both 42°C and 46°C, while MSN2/4 targets escape at 42°C. (Wilcoxon rank sum test: N.S. = P > 0.05, *** = $P$ < 0.001) (C) Colocalization was quantified by comparing the intensity of the Pab1 channel in regions with mRNA foci to the intensity of the Pab1 channel in random regions in each cell. The colocalization score is calculated separately per cell. (Wilcoxon rank sum test, N.S.: $P \geq 0.05$; **: $P < 0.01$; ***: $P < 0.001$)

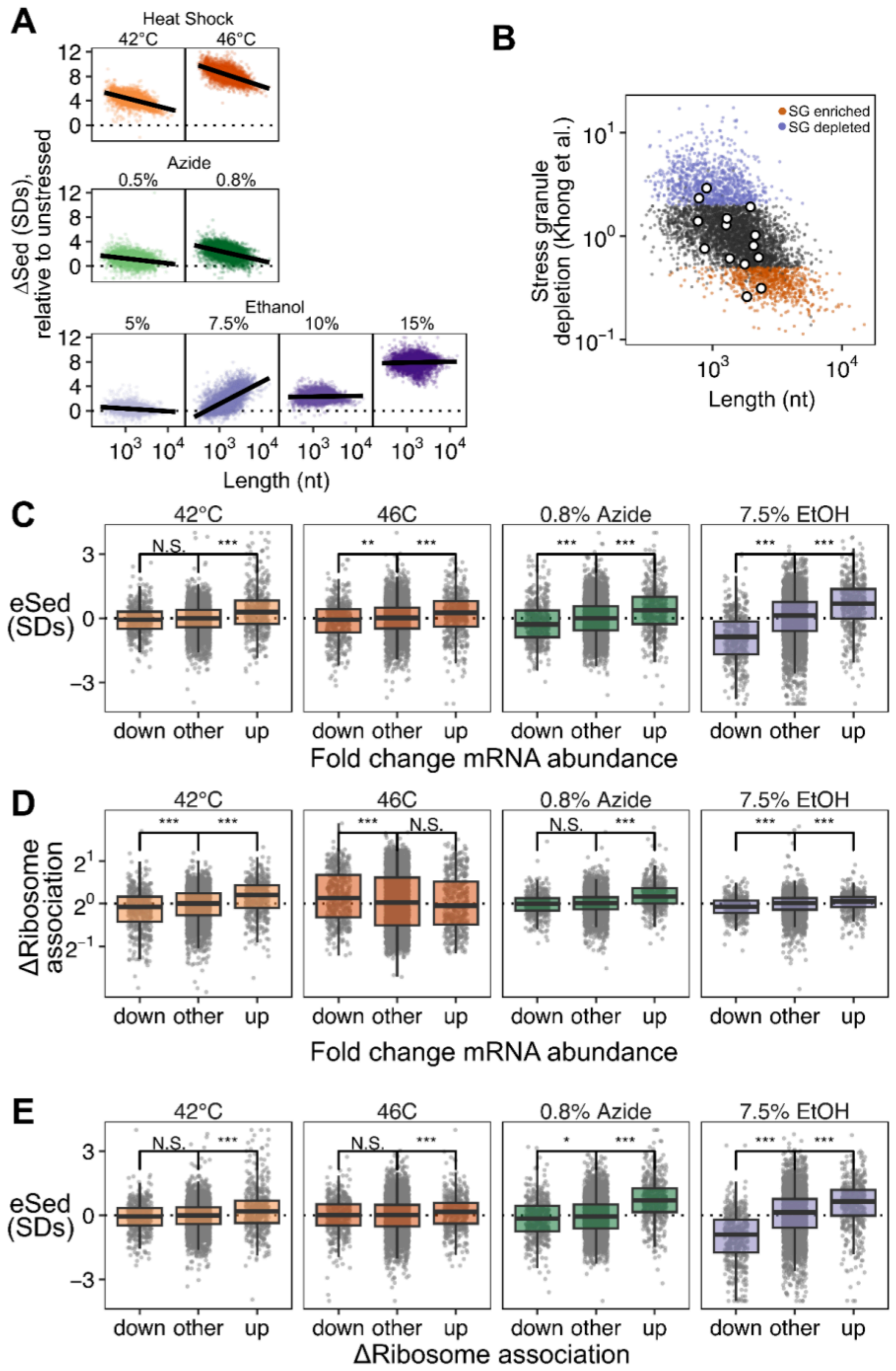

**Figure S3**: (A) Length dependence of the change in sedimentation compared across stresses. In general, shorter transcripts increase in sedimentation more than long ones. (B) Induced transcripts are not particularly depleted from stress granules based on data from Khong *et al.*, 2017. Points in white are the top 1% of induced transcripts during azide stress. (C) Comparison of escape from sedimentation (eSed) grouped by the 10% most induced and repressed transcripts during stress. (D) Comparison of the change in ribosome association grouped by the 10% most induced and repressed transcripts during stress. (E) Comparison of eSed grouped by the top and bottom 10% transcripts with changing ribosome association during stress. (Wilcoxon rank sum with Bonferroni correction: N.S. = $P \geq 0.05$, ** = $P < 0.01$, *** = $P < 0.001$)

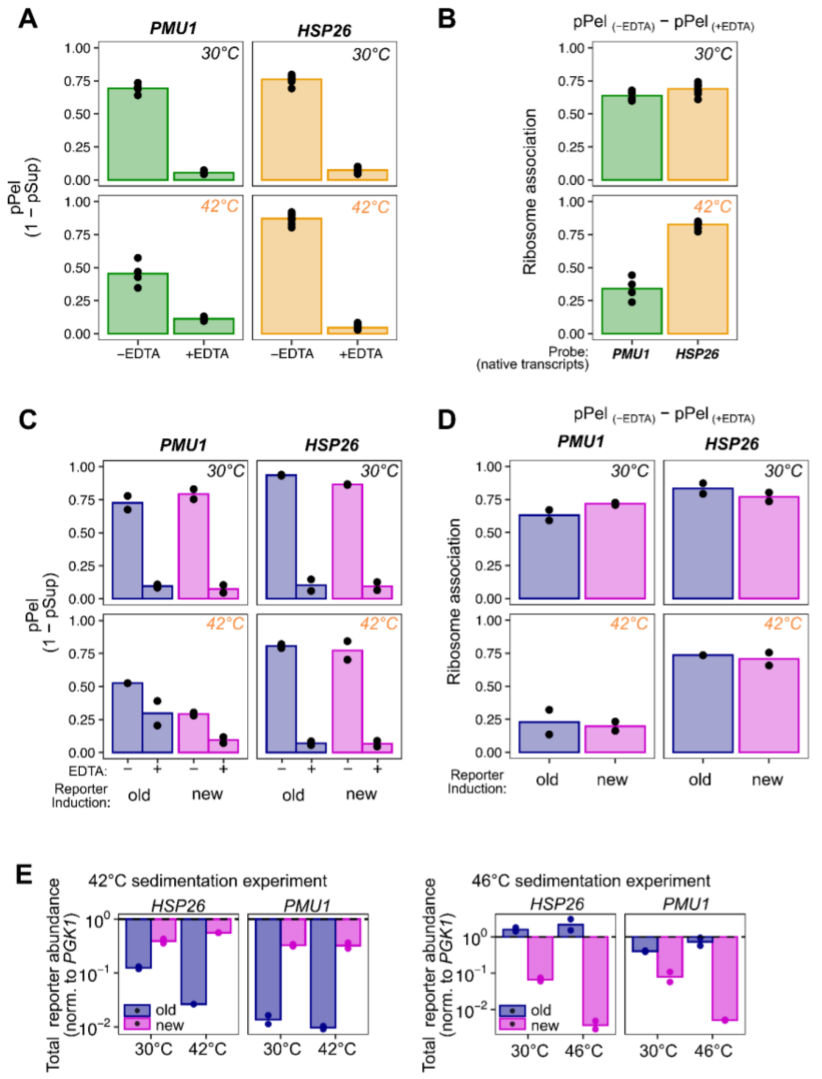

**Figure S4**: (A) Cellular lysate from stressed (20 minutes at 42°C) or unstressed yeast was centrifuged through a 1M sucrose cushion with and without EDTA. The proportion of mRNA transcripts in the pellet was quantified with qPCR, normalized to spike-in RNA. (B) In the absence of EDTA, transcripts can pellet either because they are bound to ribosomes or because they are condensed, however ribosomes fall off transcripts in the presence of EDTA. Thus to calculate the ribosome occupancy, the pPel (+EDTA) is subtracted from the pPel (−EDTA). (C,D) The same samples as in (A) and (B), but tracking the abundance of the *HSP26* and *PMU1* reporters induced before stress (old) or after stress (new). (E) Total abundance of the reporter mRNAs, measured by qPCR and normalized to native *PGK1* abundance.

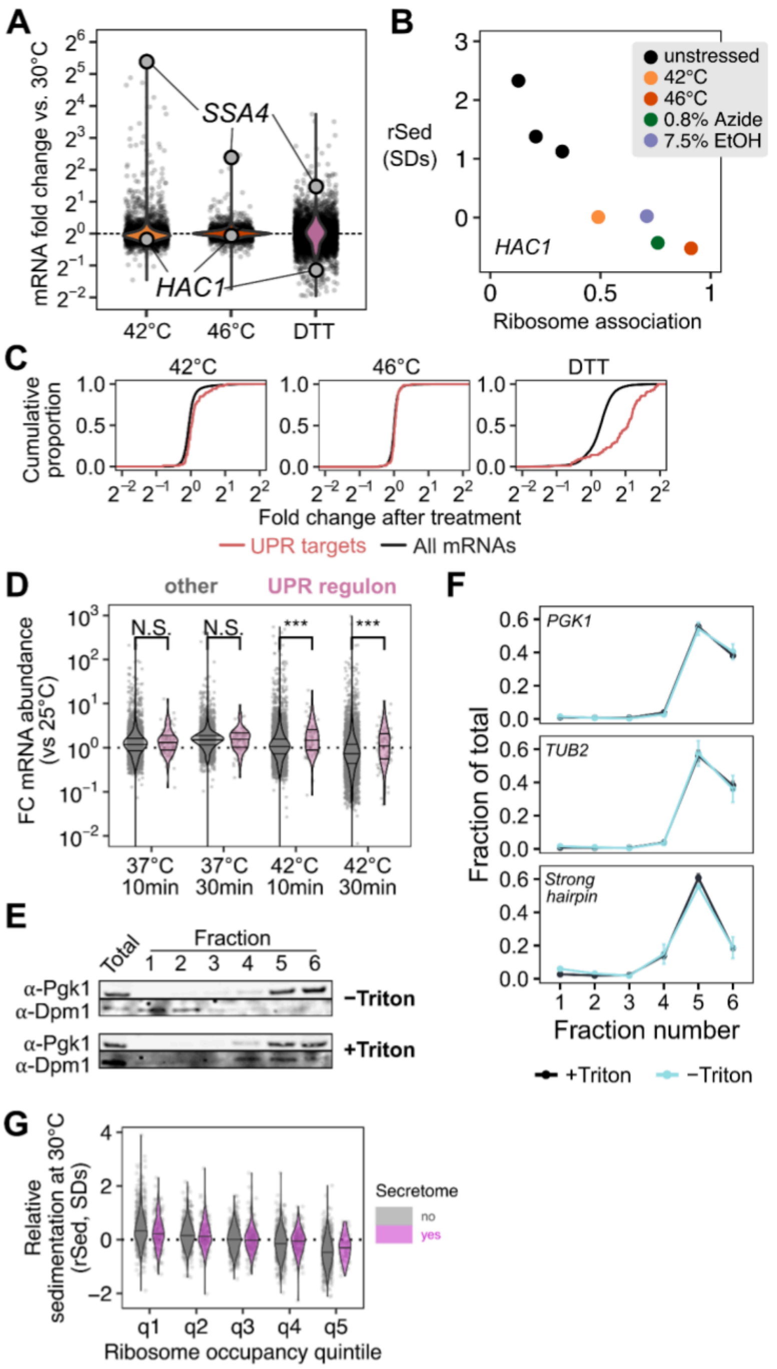

**Figure S5**: (A) *HAC1* is not induced during temperature or DTT stress. (B) *Hac1* mRNA increases in ribosome association and correspondingly decreases in relative sedimentation during stress. (C) UPR targets are slightly upregulated at 42°C and strongly upregulated in response to DTT treatment. (D) Reanalysis of Muhlhofer *et al*. 2020 RNA-seq data confirms that the UPR is upregulated after a 42°C stress but not a 37°C stress. (E) In a membrane pelleting assay, Triton X-100 disrupts the interaction between membranes and normally membrane-bound Dpm1 as measured by western blot across the fractions. (F) However, the transcript of the strong hairpin reporter, quantified by qPCR, is not found in the membrane-bound fraction with or without Triton X-100. (G) Relative sedimentation of transcripts at 30°C, categorized by ribosome occupancy (q1 is least, q5 is most) and whether the encoded protein is secreted or not (annotations from Costa et al. 2018).

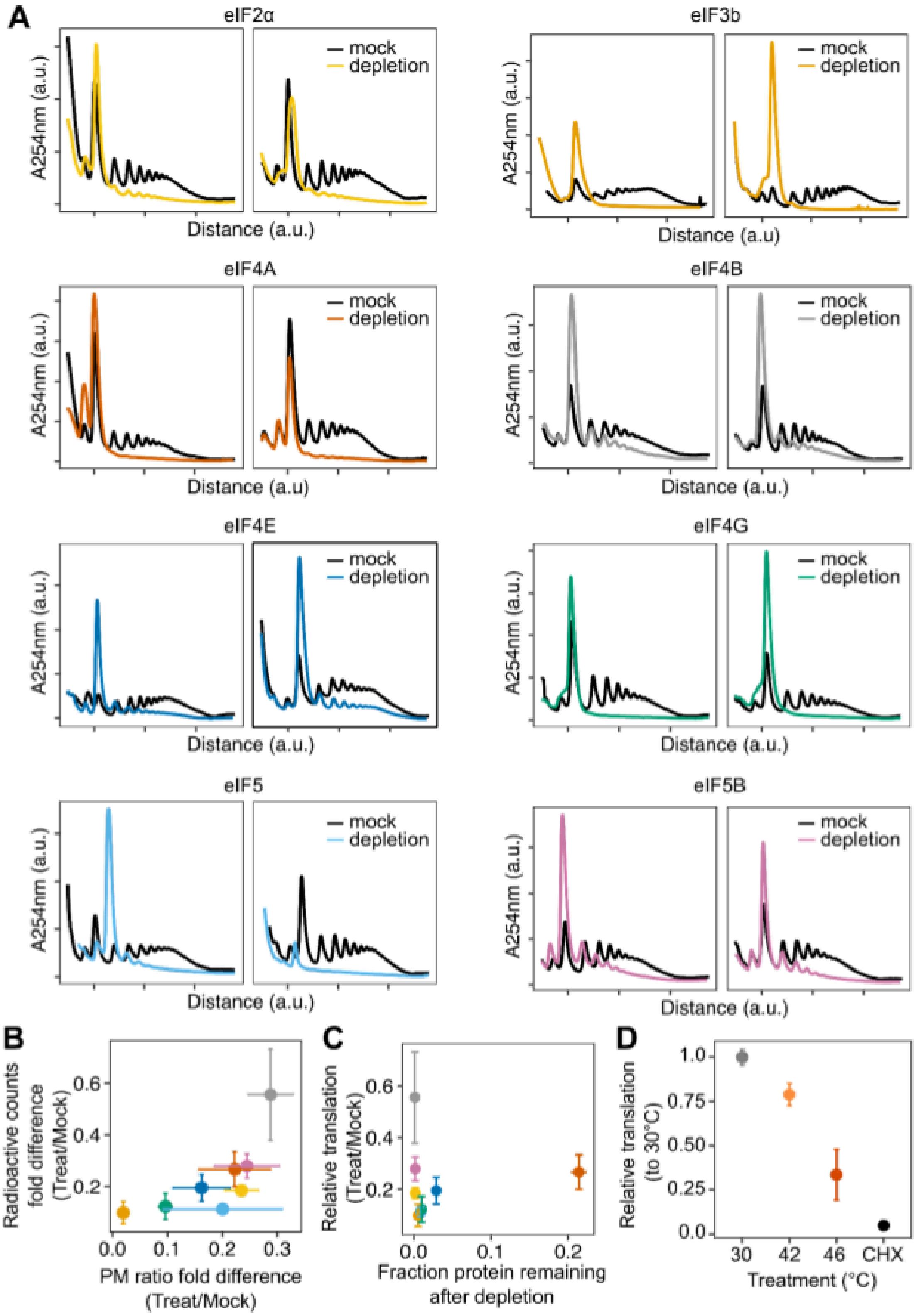

**Figure S6:** (A) Polysome profiles of AID depletion strains (B) Comparing the different measures of translation, radiolabeled amino acid incorporation and polysome profiling. (C) Comparing relative translation via radiolabelling vs. the amount of each protein remaining after the AID depletion. (D) Measures of relative translation via radiolabeled amino acid incorporation after 15 minutes of 30°C, 42°C, 46°C, or 100 µg/mL CHX.

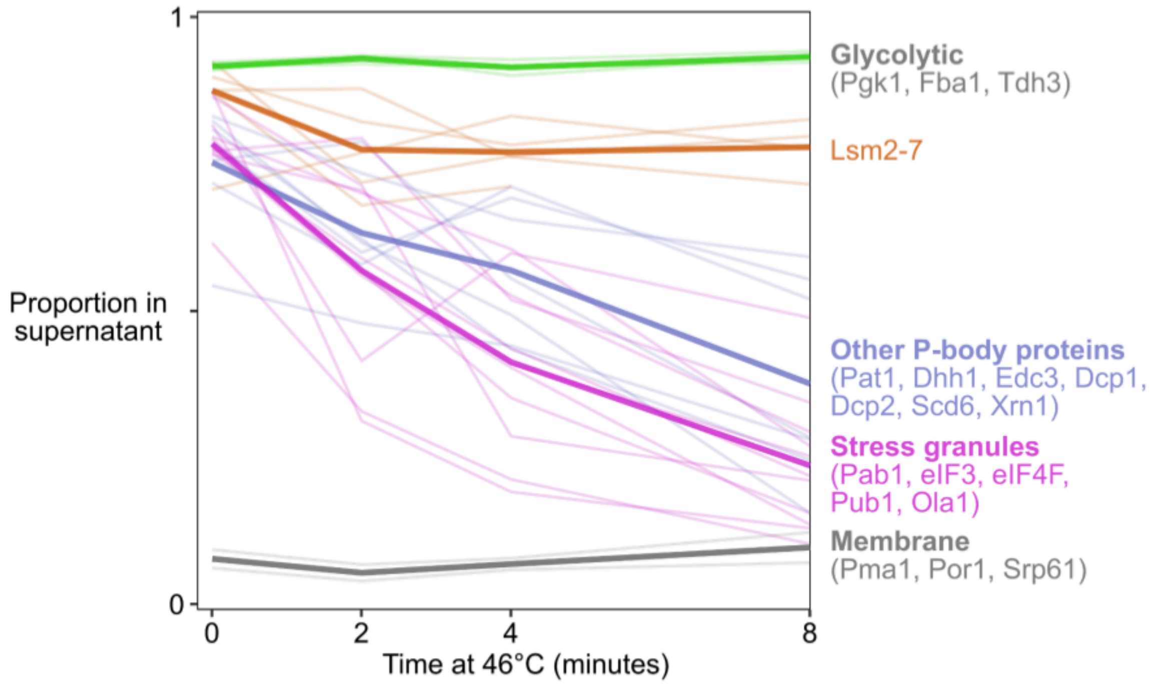

**Figure S7**: Stress induces condensation of proteins during heat shock, measured by sedimentation (data from Wallace *et al*. 2015). Under these conditions, TIICs and stress granules form. Core P-body proteins Lsm2–7 (Lsm1 is not detected) remain soluble during heat shock, while stress granule proteins and multiple other P-body proteins condense.

**Table S1**: Yeast strains used in this study

| Yeast strain | Description | Genotype | Source |
|---|---|---|---|
| BY4741 | background strain S288C | MATa ura3Δ0 leu2Δ0 his3Δ1 met15Δ0 | (Brachmann et al. 1998) |
| BY4742 | background strain S288C | MATα ura3Δ0 leu2Δ0 his3Δ1 lys2Δ0 | (Brachmann et al. 1998) |
| yJB001 | Pab1-HaloTag | MATa ura3Δ0 leu2Δ0 his3Δ1 met15Δ0 pab1::PAB1-HaloTag | This manuscript |
| yAER020 | Pab1-Clover | MATα ura3Δ0 leu2Δ0 his3Δ1 lys2Δ0 pab1::PAB1-Clover pTEF-KanMX | (Wallace et al. 2015) |
| yJB42 (eIF3b) | PRT1-AID*-3xflag+pZTRL | MATα ura3Δ0 leu2::pZ4EV-OsTIR1_Z4EV-ATF_LEU2 his3Δ1 lys2Δ0 PRT1::PRT1-AID*-3xflag | This manuscript |
| yHG55 (eIF4b) | AID*-TIF3-3xflag+pZTRL | MATα ura3Δ0 leu2::pZ4EV-OsTIR1_Z4EV-ATF_LEU2 his3Δ1 lys2Δ0 TIF3::TIF3-AID*-3xflag | This manuscript |
| yHG38 (eIF4E) | CDC33-AID*-3xflag+pZTRL | MATα ura3Δ0 leu2::pZ4EV-OsTIR1_Z4EV-ATF_LEU2 his3Δ1 lys2Δ0 CDC33::CDC33-AID*-3xflag | This manuscript |
| yHG72 (eIF4A) | TIF1-AID*-3xflag+pZTRL | MATα ura3Δ0 leu2::pZ4EV-OsTIR1_Z4EV-ATF_LEU2 his3Δ1 lys2Δ0 TIF1::TIF1-AID*-3xflag tif2Δ::KANMX | This manuscript |
| yJB143 (eIF2α) | SUI2-AID-3xFlag + pJB773 | MATα ura3Δ0 leu2::pZ4EV-OsTIR1F74G_Z4EV-ATF_LEU2 his3Δ1 lys2Δ0 SUI2::SUI2-AID*-3xflag | This manuscript |
| yJB147 (eIf5) | TIF5-AID-3xFlag + pJB773 | MATα ura3Δ0 leu2::pZ4EV-OsTIR1F74G_Z4EV-ATF_LEU2 his3Δ1 lys2Δ0 TIF5::TIF5-AID*-3xflag | This manuscript |
| yJB148 (eIF5B) | AID*-FUN12-3xFlag + pJB773 | MATα ura3Δ0 leu2::pZ4EV-OsTIR1F74G_Z4EV-ATF_LEU2 his3Δ1 lys2Δ0 FUN12::AID-FUN12*-3xflag | This manuscript |
| yJB262 (eIF4G) | TIF4631-AID*-3xFLAG+pJB773 | MATα ura3Δ0 leu2::pZ4EV-OsTIR1F74G_Z4EV-ATF_LEU2 his3Δ1 lys2Δ0 | This manuscript |

| | | | |
|---|---|---|---|
| | | TIF4631::TIF4631-AID*-3xFLAG tif4632Δ::KANMX | |
| Pbp1-GFP | Pbp1-GFP | MATa his3Δ1 leu2Δ0 met15Δ0 ura3Δ0 pbp1:: Pbp1-GFP | (Huh et al. 2003) |
| yJB265 | yJB42+Pab1-Clover | MATα ura3Δ0 leu2::pZ4EV-OsTIR1_Z4EV-ATF_LEU2 his3Δ1 lys2Δ0 PRT1::PRT1-AID*-3xflag pab1::PAB1-Clover | This manuscript |
| yHG005 | Weakest hairpin | MATa ura3Δ0 his3Δ1 met15Δ0 leu2::LEU2_5′weakest hairpin-cds2xClover-3′TPI1_5′TPI1-cdsmCherry-3′TPI1 | This manuscript |
| yHG006 | Weak hairpin | MATa ura3Δ0 his3Δ1 met15Δ0 leu2::LEU2_5′weak hairpin-cds2xClover-3′TPI1_5′TPI1-cdsmCherry-3′TPI1 | This manuscript |
| yHG007 | Medium harpin | MATa ura3Δ0 his3Δ1 met15Δ0 leu2::LEU2_5′medium hairpin-cds2xClover-3′TPI1_5′TPI1-cdsmCherry-3′TPI1 | This manuscript |
| yHG008 | Strongest hairpin | MATa ura3Δ0 his3Δ1 met15Δ0 leu2::LEU2_5′strongest hairpin-cds2xClover-3′TPI1_5′TPI1-cdsmCherry-3′TPI1 | This manuscript |
| yHG010 | No hairpin | MATa ura3Δ0 his3Δ1 met15Δ0 leu2::LEU2_5′no hairpin-cds2xClover-3′TPI1_5′TPI1-cdsmCherry-3′TPI1 | This manuscript |
| yHG026 | GCN4 reporter | MATa ura3Δ0 his3Δ1 met15Δ0 leu2::LEU2_5′GCN4-cds2xClover-3′TPI1_5′TPI1-cdsmCherry-3′TPI1 | This manuscript |
| yHG027 | GCN4 5xmut | MATa ura3Δ0 his3Δ1 met15Δ0 leu2::LEU2_5′GCN4(5xmut)-cds2xClover-3′TPI1_5′TPI1-cdsmCherry-3′TPI1 | This manuscript |
| yJB236 | Tet inducible PMU1 reporter | MATα ura3Δ0 leu2::pZ4EV-OsTIR1_Z4EV-ATF_LEU2 his3::TetR,TetR-Tup1,HIS3MX lys2Δ0 SUI2::SUI2-AID*-3xflag ho::pTETO-5′PMU1-cdsPMU1-nanoluc-3′PMU1,TPI-firefly,hygR | This manuscript |
| yJB240 | Tet inducible HSP26 | MATα ura3Δ0 | This manuscript |

| | | |
|---|---|---|
| | reporter | leu2::pZ4EV-OsTIR1_Z4EV-ATF_LEU2 his3::TetR,TetR-Tup1,HIS3MX lys2Δ0 SUI2::SUI2-AID*-3xflag ho::pTETO-5′HSP26-cdsTPI1-nanoluc-3′HSP26,TPI-firefly,hygR |

**Table S2**: Plasmids used in this study

| Plasmid | Description | Source |
|---|---|---|
| pV1382 | Cas9 editing | (Vyas et al. 2018) |
| pZTRL | Inducible OsTIR1 | (Mendoza-Ochoa et al. 2019) |
| pJB773 | OsTIR1F74G leu2 int leu | This manuscript |
| FRP2371 | TETO system | (Azizoglu, Brent, and Rudolf 2021) |
| pyHG005 | Weakest hairpin | This manuscript |
| pyHG006 | Weak hairpin | This manuscript |
| pyHG007 | Medium harpin | This manuscript |
| pyHG008 | Strongest hairpin | This manuscript |
| pyHG010 | No hairpin | This manuscript |
| pyHG026 | GCN4 reporter | This manuscript |
| pyHG027 | GCN4 5xmut | This manuscript |
| pJB801 | Tet inducible PMU1 reporter | This manuscript |
| pJB805 | Tet inducible HSP26 reporter | This manuscript |

**Table S3**: smFISH probes used in this study

| Gene | Number | Sequence | Dye |
|---|---|---|---|
| SSA4 | 1 | cacatgaataggttgtacct | Quasar 670 |
| SSA4 | 2 | tcaaccctatcgtttgcaaa | Quasar 670 |
| SSA4 | 3 | acataagaaggcgtcgttct | Quasar 670 |
| SSA4 | 4 | agcctttctgtgtcagtaaa | Quasar 670 |

| | | | |
|---|---|---|---|
| SSA4 | 5 | tattatgtgggttcatcgca | Quasar 670 |
| SSA4 | 6 | ggatcatcgaatttacgtcc | Quasar 670 |
| SSA4 | 7 | taatgcttagcatcgttcgt | Quasar 670 |
| SSA4 | 8 | cccttgtcaatcactttgaa | Quasar 670 |
| SSA4 | 9 | tctttgtctcgcctttatat | Quasar 670 |
| SSA4 | 10 | ataggctggaaccgttacta | Quasar 670 |
| SSA4 | 11 | gattgtaccggcatcttttg | Quasar 670 |
| SSA4 | 12 | tacgaagaacgttcaagccc | Quasar 670 |
| SSA4 | 13 | gcagctgtaggttcattaat | Quasar 670 |
| SSA4 | 14 | ttctgcgatttcttgtccag | Quasar 670 |
| SSA4 | 15 | aaagatcaagacgttgtgct | Quasar 670 |
| SSA4 | 16 | ccttcatctatggatagcag | Quasar 670 |
| SSA4 | 17 | agaaagttaaccagcctact | Quasar 670 |
| SSA4 | 18 | ttttcttttgaactcctcgg | Quasar 670 |
| SSA4 | 19 | ggttagttgttagatccttt | Quasar 670 |
| SSA4 | 20 | tccttaacctccttagggac | Quasar 670 |
| SSA4 | 21 | tttctatagatgtctgagca | Quasar 670 |
| SSA4 | 22 | aattcttcaaatcttgccct | Quasar 670 |
| SSA4 | 23 | atcagccaaaactttttcca | Quasar 670 |
| SSA4 | 24 | accagtttttgtacttttgg | Quasar 670 |
| SSA4 | 25 | cctcatcagggttaatcgaa | Quasar 670 |
| SSA4 | 26 | ctgtacggcagcaccataag | Quasar 670 |
| SSA4 | 27 | actggtcacccgttaagatg | Quasar 670 |
| SSA4 | 28 | cagtaaatcttgggtcgtcg | Quasar 670 |
| SSA4 | 29 | ataatggtgcaacatccagc | Quasar 670 |
| SSA4 | 30 | ttttttgttgggatagtcga | Quasar 670 |
| SSA4 | 31 | gtaggtggaaaacacttccg | Quasar 670 |
| SSA4 | 32 | tttcaccctcaaaaacttgt | Quasar 670 |
| SSA4 | 33 | caactcaaatttacccagta | Quasar 670 |
| SSA4 | 34 | tcaacggcagatacgttcag | Quasar 670 |
| SSA4 | 35 | tcttcccttatcgttagtaa | Quasar 670 |
| SSA4 | 36 | aacttttctgcctcagcaac | Quasar 670 |
| SSA4 | 37 | gattcttagcttgaacacgt | Quasar 670 |
| SSA4 | 38 | gtaaacgcgtacgattctag | Quasar 670 |
| SSA4 | 39 | ttcgctcacagaatttttca | Quasar 670 |

| | | | |
|---|---|---|---|
| SSA4 | 40 | ccaccttctccttgaagtta | Quasar 670 |
| SSA4 | 41 | aatttcctggcatcctcttc | Quasar 670 |
| SSA4 | 42 | atttatagcatcttgggcgg | Quasar 670 |
| SSA4 | 43 | ccgcttgcgaagcatctaac | Quasar 670 |
| SSA4 | 44 | ataatggggtttgcaacacc | Quasar 670 |
| SSA4 | 45 | ctgcagctccgtaaaattta | Quasar 670 |
| SSB1 | 1 | taccgatagcaccttggaaa | Quasar 670 |
| SSB1 | 2 | tttcttgggttcaaagcagc | Quasar 670 |
| SSB1 | 3 | gtcgtcgaatcttctaccaa | Quasar 670 |
| SSB1 | 4 | gtcgataaccttgaaaggcc | Quasar 670 |
| SSB1 | 5 | tcttggtttcttccaagtat | Quasar 670 |
| SSB1 | 6 | agcggaaatttcttgtgggg | Quasar 670 |
| SSB1 | 7 | accaatcttagcttcagcaa | Quasar 670 |
| SSB1 | 8 | ggttcgttgatgatacgcaa | Quasar 670 |
| SSB1 | 9 | cacctagaccgtaagcaata | Quasar 670 |
| SSB1 | 10 | aacatgtctttccttttcgg | Quasar 670 |
| SSB1 | 11 | accagcaatgtgcaacaagg | Quasar 670 |
| SSB1 | 12 | gtgttaccggaagtagattt | Quasar 670 |
| SSB1 | 13 | agtgttccaacaagttggtg | Quasar 670 |
| SSB1 | 14 | ttcttcttgaattcagcctt | Quasar 670 |
| SSB1 | 15 | atcgtcggagatgtccaaac | Quasar 670 |
| SSB1 | 16 | aaggttctcttagctctttc | Quasar 670 |
| SSB1 | 17 | acggtagtttgagtgacaga | Quasar 670 |
| SSB1 | 18 | ggattcgaaatcttcaccgt | Quasar 670 |
| SSB1 | 19 | aacaatgcggcgttcaagtc | Quasar 670 |
| SSB1 | 20 | cttagagatcttagcatcct | Quasar 670 |
| SSB1 | 21 | ccaaccaagacaacttcgtc | Quasar 670 |
| SSB1 | 22 | tccaattgcttaccgtcaaa | Quasar 670 |
| SSB1 | 23 | acaacaagtccttggtttcg | Quasar 670 |
| SSB1 | 24 | gttcttctcttgatggttgg | Quasar 670 |
| SSB1 | 25 | actgggaattgaacggtggt | Quasar 670 |
| SSB1 | 26 | tgggatgttcttcaagtcga | Quasar 670 |
| SSB1 | 27 | atagcttccaagactggttc | Quasar 670 |

| | | | |
|---|---|---|---|
| SSB1 | 28 | accgttagcatcaacttcga | Quasar 670 |
| SSB1 | 29 | cgacggcagtaaccttcaag | Quasar 670 |
| SSB1 | 30 | gaagacttaccggtagactt | Quasar 670 |
| SSB1 | 31 | agacaagactgggtcagtga | Quasar 670 |
| SSB1 | 32 | gacttggaacctctcttcaa | Quasar 670 |
| SSB1 | 33 | tcgatttgcaaagcagccaa | Quasar 670 |
| SSB1 | 34 | aacgagaagacatggccttg | Quasar 670 |
| HSP104 | 1 | taggtaagggactgatccat | Quasar 670 |
| HSP104 | 2 | caggttgctgttgaggaatt | Quasar 670 |
| HSP104 | 3 | cccaaagcataacttggagt | Quasar 670 |
| HSP104 | 4 | ttgaatcttagcagcgtctt | Quasar 670 |
| HSP104 | 5 | gcgctataaatgagtccttc | Quasar 670 |
| HSP104 | 6 | tggcctcaatatctacttga | Quasar 670 |
| HSP104 | 7 | accacgaagttcaagagctt | Quasar 670 |
| HSP104 | 8 | ccacgagagtcaattctagt | Quasar 670 |
| HSP104 | 9 | tccaaaggtgtgttcgtatc | Quasar 670 |
| HSP104 | 10 | cctgctcagtcatatcaatg | Quasar 670 |
| HSP104 | 11 | agggtcaagtttaccttgac | Quasar 670 |
| HSP104 | 12 | tcttatttcttcttcacggc | Quasar 670 |
| HSP104 | 13 | acctggctcaccaattaaac | Quasar 670 |
| HSP104 | 14 | ttagcgccttgtaagatagt | Quasar 670 |
| HSP104 | 15 | aatgcggccaaatctagact | Quasar 670 |
| HSP104 | 16 | cttcaagatgttagcagcgt | Quasar 670 |
| HSP104 | 17 | tcaattggcctctggacaaa | Quasar 670 |
| HSP104 | 18 | cttctttcaaaggcaccatc | Quasar 670 |
| HSP104 | 19 | cactgtttgtctcacacttg | Quasar 670 |
| HSP104 | 20 | gttgcagacctctcaatatg | Quasar 670 |
| HSP104 | 21 | actaaggcgctatccagaat | Quasar 670 |
| HSP104 | 22 | aacgcttggctaattgagca | Quasar 670 |
| HSP104 | 23 | ctggcaatcttctatatggc | Quasar 670 |
| HSP104 | 24 | tcttctggcttagaatctct | Quasar 670 |
| HSP104 | 25 | catcttcatctctctctaga | Quasar 670 |
| HSP104 | 26 | ctatctttagtggtggagtc | Quasar 670 |

| | | | |
|---|---|---|---|
| HSP104 | 27 | tgcaatgaagcttccttctg | Quasar 670 |
| HSP104 | 28 | gttgtcttagaggttccaat | Quasar 670 |
| HSP104 | 29 | tttgatatctgggatggcga | Quasar 670 |
| HSP104 | 30 | accacattttggatcatgga | Quasar 670 |
| HSP104 | 31 | atcttgcagctgtttcagaa | Quasar 670 |
| HSP104 | 32 | cccacgacttcagatgataa | Quasar 670 |
| HSP104 | 33 | acggcattggaaacagcttt | Quasar 670 |
| HSP104 | 34 | ccttggattagctaaacctg | Quasar 670 |
| HSP104 | 35 | tagccaattcagttttaccg | Quasar 670 |
| HSP104 | 36 | aatcgaccctgatcatcatg | Quasar 670 |
| HSP104 | 37 | acttagagaccgcatacttc | Quasar 670 |
| HSP104 | 38 | ttcatcgtacccgacataac | Quasar 670 |
| HSP104 | 39 | cggagtatggtttgtattgc | Quasar 670 |
| HSP104 | 40 | gtaattctaccgtcatccaa | Quasar 670 |
| HSP104 | 41 | ttggaacagtcgatcgtctt | Quasar 670 |
| HSP104 | 42 | aaatgttgcctaacagcacc | Quasar 670 |
| HSP104 | 43 | tttgctcgaatctctcttca | Quasar 670 |
| HSP104 | 44 | ggcctcttgagttaaattca | Quasar 670 |
| HSP104 | 45 | cccatatcatcggaataacc | Quasar 670 |
| HSP104 | 46 | tctttaagatccttagtgcc | Quasar 670 |
| HSP104 | 47 | cgagatttacccttcttcaa | Quasar 670 |
| HSP104 | 48 | tagtagcttcgtgatttggt | Quasar 670 |
| ADD66 | 1 | gtatgttcctttgggttttg | Quasar 670 |
| ADD66 | 2 | tactaatggcaacaccaggc | Quasar 670 |
| ADD66 | 3 | attgtggtatattccctacg | Quasar 670 |
| ADD66 | 4 | ttcagcagccagtcaatact | Quasar 670 |
| ADD66 | 5 | ttcccattcatttgcttgtg | Quasar 670 |
| ADD66 | 6 | tcgaatctaacgcctccaaa | Quasar 670 |
| ADD66 | 7 | cccacaaattccactaggta | Quasar 670 |
| ADD66 | 8 | atcctctggtctgtctaatg | Quasar 670 |
| ADD66 | 9 | tctttatatagcgagtcgct | Quasar 670 |
| ADD66 | 10 | caaagcgctgctatacttca | Quasar 670 |
| ADD66 | 11 | cctcgcttcttattgtagaa | Quasar 670 |

| | | | |
|---|---|---|---|
| ADD66 | 12 | ttcgttgctgtatggcaaaa | Quasar 670 |
| ADD66 | 13 | tagttaacggacacgagtgg | Quasar 670 |
| ADD66 | 14 | cggcaagatgatctctacga | Quasar 670 |
| ADD66 | 15 | ccgatatattatacttgctc | Quasar 670 |
| ADD66 | 16 | agagagtcccaaatgcatat | Quasar 670 |
| ADD66 | 17 | attttcatcctccattgcat | Quasar 670 |
| ADD66 | 18 | cctgtggacgtacaatcaca | Quasar 670 |
| ADD66 | 19 | aactctccgagcgaatatac | Quasar 670 |
| ADD66 | 20 | ctcagcttcatcgtcgaaat | Quasar 670 |
| ADD66 | 21 | tcgtttagatggaggttcga | Quasar 670 |
| ADD66 | 22 | gttattgaccatcgactctt | Quasar 670 |
| ADD66 | 23 | tcttggaaactggtaggagt | Quasar 670 |
| ADD66 | 24 | tcggctgatcaacggatatc | Quasar 670 |
| ADD66 | 25 | cgacgcattcagtatttgga | Quasar 670 |
| ADD66 | 26 | taatgctgcggagggcttta | Quasar 670 |
| ADD66 | 27 | attggccaaacaactacagt | Quasar 670 |
| ADD66 | 28 | gaatctaatgagttgtcccc | Quasar 670 |
| ADD66 | 29 | ggcgcgtttttgataacttt | Quasar 670 |
| ADD66 | 30 | ggcctgacaaactttacaat | Quasar 670 |
| ADD66 | 31 | gtatgcaccttgccatgata | Quasar 670 |
| ADD66 | 32 | aatttatctcttgcatccgc | Quasar 670 |
| ADD66 | 33 | tacttctcggtttcaattgt | Quasar 670 |
| ADD66 | 34 | gtggctatatatgcacttgt | Quasar 670 |
| oligodT | 1 | tttttttttttttttttttt | Quasar 570 |

**Table S4:** Sequencing information:

| Lysate_ sample | Temperature | Biorep | Strain | Treatment Group | Treatment | Experiment | Sequencing Group (see Methods) | spike-in |
|---|---|---|---|---|---|---|---|---|
| L115 | 30C | BR1 | BY4741 | PolySeq_0.8% Azide | Azide_0.8 %_pH6.8 | PolySeq | C | pombe |
| L117 | 30C | BR2 | BY4741 | PolySeq_0.8% Azide | Azide_0.8 %_pH6.8 | PolySeq | C | pombe |

| | | | | | | | | |
|---|---|---|---|---|---|---|---|---|
| L114 | 30C | BR1 | BY4741 | PolySeq_0.8% Azide | mock | PolySeq | C | pombe |
| L116 | 30C | BR2 | BY4741 | PolySeq_0.8% Azide | mock | PolySeq | C | pombe |
| L51 | 30C | BR1 | BY4741 | PolySeq_10min | none | PolySeq | B | pombe |
| L54 | 30C | BR2 | BY4741 | PolySeq_10min | none | PolySeq | B | pombe |
| L52 | 42C | BR1 | BY4741 | PolySeq_10min | none | PolySeq | B | pombe |
| L55 | 42C | BR2 | BY4741 | PolySeq_10min | none | PolySeq | B | pombe |
| L53 | 46C | BR1 | BY4741 | PolySeq_10min | none | PolySeq | B | pombe |
| L56 | 46C | BR2 | BY4741 | PolySeq_10min | none | PolySeq | B | pombe |
| L123 | 30C | BR1 | BY4741 | PolySeq_7.5% EtOH | EtOH_7.5 % | PolySeq | C | pombe |
| L125 | 30C | BR2 | BY4741 | PolySeq_7.5% EtOH | EtOH_7.5 % | PolySeq | C | pombe |
| L122 | 30C | BR1 | BY4741 | PolySeq_7.5% EtOH | mock | PolySeq | C | pombe |
| L124 | 30C | BR2 | BY4741 | PolySeq_7.5% EtOH | mock | PolySeq | C | pombe |
| L82 | 30C | BR1 | BY4741 | Azide_0.5%_p H6.8 | Azide_0.5 %_pH6.8 | SedSeq | B | nanoluc |
| L84 | 30C | BR2 | BY4741 | Azide_0.5%_p H6.8 | Azide_0.5 %_pH6.8 | SedSeq | B | nanoluc |
| L81 | 30C | BR1 | BY4741 | Azide_0.5%_p H6.8 | mock | SedSeq | B | nanoluc |
| L83 | 30C | BR2 | BY4741 | Azide_0.5%_p H6.8 | mock | SedSeq | B | nanoluc |
| L107 | 30C | BR1 | BY4741 | azide3 | Azide_0.8 %_pH6.8 | SedSeq | C | nanoluc |
| L109 | 30C | BR2 | BY4741 | azide3 | Azide_0.8 %_pH6.8 | SedSeq | C | nanoluc |
| L106 | 30C | BR1 | BY4741 | azide3 | mock | SedSeq | C | nanoluc |
| L108 | 30C | BR2 | BY4741 | azide3 | mock | SedSeq | C | nanoluc |
| L33 | 30C | BR1 | BY4741 | CHX | CHX | SedSeq | B | nanoluc |
| L38 | 30C | BR2 | BY4741 | CHX | CHX | SedSeq | B | nanoluc |
| L31 | 30C | BR1 | BY4741 | CHX | mock | SedSeq | B | nanoluc |
| L35 | 30C | BR2 | BY4741 | CHX | mock | SedSeq | B | nanoluc |
| L39 | 42C | BR2 | BY4741 | CHX | CHX | SedSeq | B | nanoluc |
| L36 | 42C | BR2 | BY4741 | CHX | mock | SedSeq | B | nanoluc |

| | | | | | | | | |
|---|---|---|---|---|---|---|---|---|
| L34 | 46C | BR1 | BY4741 | CHX | CHX | SedSeq | B | nanoluc |
| L40 | 46C | BR2 | BY4741 | CHX | CHX | SedSeq | B | nanoluc |
| L32 | 46C | BR1 | BY4741 | CHX | mock | SedSeq | B | nanoluc |
| L37 | 46C | BR2 | BY4741 | CHX | mock | SedSeq | B | nanoluc |
| L72 | 30C | BR1 | yJB042 | eIF3b_deplete | eIF3b_deplete | SedSeq | B | nanoluc |
| L78 | 30C | BR2 | yJB042 | eIF3b_deplete | eIF3b_deplete | SedSeq | B | nanoluc |
| L69 | 30C | BR1 | yJB042 | eIF3b_deplete | mock | SedSeq | B | nanoluc |
| L75 | 30C | BR2 | yJB042 | eIF3b_deplete | mock | SedSeq | B | nanoluc |
| L73 | 42C | BR1 | yJB042 | eIF3b_deplete | eIF3b_deplete | SedSeq | B | nanoluc |
| L79 | 42C | BR2 | yJB042 | eIF3b_deplete | eIF3b_deplete | SedSeq | B | nanoluc |
| L70 | 42C | BR1 | yJB042 | eIF3b_deplete | mock | SedSeq | B | nanoluc |
| L76 | 42C | BR2 | yJB042 | eIF3b_deplete | mock | SedSeq | B | nanoluc |
| L74 | 46C | BR1 | yJB042 | eIF3b_deplete | eIF3b_deplete | SedSeq | B | nanoluc |
| L80 | 46C | BR2 | yJB042 | eIF3b_deplete | eIF3b_deplete | SedSeq | B | nanoluc |
| L71 | 46C | BR1 | yJB042 | eIF3b_deplete | mock | SedSeq | B | nanoluc |
| L77 | 46C | BR2 | yJB042 | eIF3b_deplete | mock | SedSeq | B | nanoluc |
| L60 | 30C | BR1 | yHG038 | eIF4E_deplete | eIF4E_deplete | SedSeq | B | nanoluc |
| L66 | 30C | BR2 | yHG038 | eIF4E_deplete | eIF4E_deplete | SedSeq | B | nanoluc |
| L57 | 30C | BR1 | yHG038 | eIF4E_deplete | mock | SedSeq | B | nanoluc |
| L63 | 30C | BR2 | yHG038 | eIF4E_deplete | mock | SedSeq | B | nanoluc |
| L61 | 42C | BR1 | yHG038 | eIF4E_deplete | eIF4E_deplete | SedSeq | B | nanoluc |
| L67 | 42C | BR2 | yHG038 | eIF4E_deplete | eIF4E_deplete | SedSeq | B | nanoluc |
| L58 | 42C | BR1 | yHG038 | eIF4E_deplete | mock | SedSeq | B | nanoluc |
| L64 | 42C | BR2 | yHG038 | eIF4E_deplete | mock | SedSeq | B | nanoluc |
| L62 | 46C | BR1 | yHG038 | eIF4E_deplete | eIF4E_deplete | SedSeq | B | nanoluc |
| L68 | 46C | BR2 | yHG038 | eIF4E_deplete | eIF4E_deplete | SedSeq | B | nanoluc |

| | | | | | | | | |
|---|---|---|---|---|---|---|---|---|
| L59 | 46C | BR1 | yHG038 | eIF4E_deplete | mock | SedSeq | B | nanoluc |
| L65 | 46C | BR2 | yHG038 | eIF4E_deplete | mock | SedSeq | B | nanoluc |
| L7 | 30C | BR1 | BY4741 | EW_TSPP | none | SedSeq | A | none |
| L11 | 30C | BR2 | BY4741 | EW_TSPP | none | SedSeq | A | none |
| L8 | 42C | BR1 | BY4741 | EW_TSPP | none | SedSeq | A | none |
| L12 | 42C | BR2 | BY4741 | EW_TSPP | none | SedSeq | A | none |
| L9 | 46C | BR1 | BY4741 | EW_TSPP | none | SedSeq | A | none |
| L13 | 46C | BR2 | BY4741 | EW_TSPP | none | SedSeq | A | none |
| L89 | 30C | BR1 | yHG010 | hairpinReporters | none | SedSeq | C | nanoluc |
| L90 | 30C | BR1 | yHG005 | hairpinReporters | none | SedSeq | C | nanoluc |
| L91 | 30C | BR1 | yHG006 | hairpinReporters | none | SedSeq | C | nanoluc |
| L92 | 30C | BR1 | yHG007 | hairpinReporters | none | SedSeq | C | nanoluc |
| L93 | 30C | BR1 | yHG008 | hairpinReporters | none | SedSeq | C | nanoluc |
| L94 | 30C | BR1 | yHG026 | hairpinReporters | none | SedSeq | C | nanoluc |
| L95 | 30C | BR1 | yHG027 | hairpinReporters | none | SedSeq | C | nanoluc |
| L43 | 30C | BR1 | BY4741 | HyperOsmotic_EtOH | EtOH_15% | SedSeq | B | nanoluc |
| L46 | 30C | BR2 | BY4741 | HyperOsmotic_EtOH | EtOH_15% | SedSeq | B | nanoluc |
| L41 | 30C | BR1 | BY4741 | HyperOsmotic_EtOH | mock | SedSeq | B | nanoluc |
| L44 | 30C | BR2 | BY4741 | HyperOsmotic_EtOH | mock | SedSeq | B | nanoluc |
| L98 | 30C | BR1 | BY4741 | milderEtOH1 | EtOH_10% | SedSeq | C | nanoluc |
| L101 | 30C | BR2 | BY4741 | milderEtOH1 | EtOH_10% | SedSeq | C | nanoluc |
| L97 | 30C | BR1 | BY4741 | milderEtOH1 | EtOH_5% | SedSeq | C | nanoluc |
| L100 | 30C | BR2 | BY4741 | milderEtOH1 | EtOH_5% | SedSeq | C | nanoluc |
| L96 | 30C | BR1 | BY4741 | milderEtOH1 | mock | SedSeq | C | nanoluc |
| L99 | 30C | BR2 | BY4741 | milderEtOH1 | mock | SedSeq | C | nanoluc |

| | | | | | | | | |
|---|---|---|---|---|---|---|---|---|
| L103 | 30C | BR1 | BY4741 | milderEtOH2 | EtOH_7.5% | SedSeq | C | nanoluc |
| L105 | 30C | BR2 | BY4741 | milderEtOH2 | EtOH_7.5% | SedSeq | C | nanoluc |
| L102 | 30C | BR1 | BY4741 | milderEtOH2 | mock | SedSeq | C | nanoluc |
| L104 | 30C | BR2 | BY4741 | milderEtOH2 | mock | SedSeq | C | nanoluc |
| L110 | 30C | BR1 | BY4741 | spombe_spikeIn | none | SedSeq | C | nanoluc+pombe |
| L112 | 30C | BR2 | BY4741 | spombe_spikeIn | none | SedSeq | C | nanoluc+pombe |
| L111 | 46C | BR1 | BY4741 | spombe_spikeIn | none | SedSeq | C | nanoluc+pombe |
| L113 | 46C | BR2 | BY4741 | spombe_spikeIn | none | SedSeq | C | nanoluc+pombe |
| L49 | 30C | BR1 | BY4741 | uprInduction | DTT_10mM | SedSeq | B | nanoluc |
| L50 | 30C | BR2 | BY4741 | uprInduction | DTT_10mM | SedSeq | B | nanoluc |
| L47 | 30C | BR1 | BY4741 | uprInduction | mock | SedSeq | B | nanoluc |
| L48 | 30C | BR2 | BY4741 | uprInduction | mock | SedSeq | B | nanoluc |

Azizoglu, Asli, Roger Brent, and Fabian Rudolf. 2021. "A Precisely Adjustable, Variation-Suppressed Eukaryotic Transcriptional Controller to Enable Genetic Discovery." *eLife* 10 (August). https://doi.org/10.7554/eLife.69549.

Brachmann, C. B., A. Davies, G. J. Cost, E. Caputo, J. Li, P. Hieter, and J. D. Boeke. 1998. "Designer Deletion Strains Derived from Saccharomyces Cerevisiae S288C: A Useful Set of Strains and Plasmids for PCR-Mediated Gene Disruption and Other Applications." *Yeast* 14 (2): 115–32.

Huh, Won-Ki, James V. Falvo, Luke C. Gerke, Adam S. Carroll, Russell W. Howson, Jonathan S. Weissman, and Erin K. O'Shea. 2003. "Global Analysis of Protein Localization in Budding Yeast." *Nature* 425 (6959): 686–91.

Mendoza-Ochoa, Gonzalo I., J. David Barrass, Barbara R. Terlouw, Isabella E. Maudlin, Susana de Lucas, Emanuela Sani, Vahid Aslanzadeh, Jane A. E. Reid, and Jean D. Beggs. 2019. "A Fast and Tuneable Auxin-Inducible Degron for Depletion of Target Proteins in Budding Yeast." *Yeast* 36 (1): 75–81.

Reid, David W., and Christopher V. Nicchitta. 2015. "Diversity and Selectivity in mRNA Translation on the Endoplasmic Reticulum." *Nature Reviews. Molecular Cell Biology* 16 (4): 221–31.

Tudek, Agnieszka, Paweł S. Krawczyk, Seweryn Mroczek, Rafał Tomecki, Matti Turtola, Katarzyna Matylla-Kulińska, Torben Heick Jensen, and Andrzej Dziembowski. 2021. "Global View on the Metabolism of RNA poly(A) Tails in Yeast Saccharomyces Cerevisiae." *Nature Communications* 12 (1): 4951.

Vyas, Valmik K., G. Guy Bushkin, Douglas A. Bernstein, Matthew A. Getz, Magdalena Sewastianik, M. Inmaculada Barrasa, David P. Bartel, and Gerald R. Fink. 2018. "New

CRISPR Mutagenesis Strategies Reveal Variation in Repair Mechanisms among Fungi." *mSphere* 3 (2). https://doi.org/10.1128/mSphere.00154-18.

Wallace, Edward W. J., Jamie L. Kear-Scott, Evgeny V. Pilipenko, Michael H. Schwartz, Pawel R. Laskowski, Alexandra E. Rojek, Christopher D. Katanski, et al. 2015. "Reversible, Specific, Active Aggregates of Endogenous Proteins Assemble upon Heat Stress." *Cell* 162 (6): 1286–98.

Yoffe, Aron M., Peter Prinsen, Ajaykumar Gopal, Charles M. Knobler, William M. Gelbart, and Avinoam Ben-Shaul. 2008. "Predicting the Sizes of Large RNA Molecules." *Proceedings of the National Academy of Sciences of the United States of America* 105 (42): 16153–58.
